# gp130/STAT3 signaling is required for homeostatic proliferation and anabolism in postnatal growth plate and articular chondrocytes

**DOI:** 10.1101/2021.10.12.464120

**Authors:** Nancy Q. Liu, Yucheng Lin, Liangliang Li, Jinxiu Lu, Dawei Geng, Jiankang Zhang, Tea Jashashvili, Zorica Buser, Jenny Magallanes, Jade Tassey, Ruzanna Shkhyan, Arijita Sarkar, Noah Lopez, Liming Wang, Frank A. Petrigliano, Ben Van Handel, Karen Lyons, Denis Evseenko

## Abstract

Growth of long bones and vertebrae is maintained postnatally by a long-lasting pool of progenitor cells. Little is known about the molecular mechanisms that regulate the output and maintenance of the cells that give rise to mature cartilage. Here we demonstrate that postnatal chondrocyte-specific deletion of a transcription factor Stat3 results in severely reduced proliferation coupled with increased hypertrophy, growth plate fusion, stunting and signs of progressive dysfunction of the articular cartilage. This effect is dimorphic, with females more strongly affected than males. Chondrocyte-specific deletion of the IL-6 family cytokine receptor gp130, which activates Stat3, phenocopied Stat3^-^deletion; deletion of Lifr, one of many co-receptors that signals through gp130, resulted in a milder phenotype. These data define a new molecular circuit that regulates chondrogenic cell maintenance and output and reveals a novel, hitherto unrecognized function of IL-6 cytokines in the skeletal system with direct implications for skeletal development and regeneration.

## Introduction

The continual growth of long bones and vertebrae is fueled by chondrogenic stem and progenitor cells in the growth plate^1^. Beginning in embryogenesis and continuing through adolescence, these chondroprogenitors provide a steady stream of progeny that undergo endochondral ossification and contribute to bone growth^2^. Chondrocytes enter a phase of rapid division as they leave the resting zone of the growth plate, forming characteristic columns of mitotic cells that begin to secrete increased levels of extracellular matrix (ECM) proteins such as collagen II and aggrecan^3^. Once their distance from the niche within the resting zone increases to its maximum extent, they become hypertrophic, deposit lower amounts of ECM, and begin secreting collagen X and alkaline phosphatase, which is essential for calcification of the ECM^4^. Once surrounded by calcified matrix, these chondrocytes either undergo apoptosis or transdifferentiate into osteoprogenitors^5^; osteoprogenitors derived from the growth plate, or invading from the periosteum, differentiate into osteoblasts and secrete bone matrix. Endochondral ossification takes place during fracture repair, with similar coordinated cellular dynamics^6^.

In humans, growth plates become inactive at the end of puberty, resulting in replacement with bone; this occurs earlier in the long bones than in the spine^7^. Females undergo growth plate fusion an average of 2 years earlier than males, suggesting sexually dimorphic regulation of chondroprogenitors^8^. In mice, growth plates in the long bones and spine persist throughout the lifespan of the animal, providing an opportunity to study how specific molecular pathways impact the dynamics of chondroprogenitors. Recently, Newton et al. defined a population of chondrogenic cells localized to the growth plate that generate clonal progeny and support long bone growth^9^. Their data supported previous work showing that Hedgehog (Hh) signaling is required for growth plate maintenance; if this pathway is inhibited, growth plates fuse and differentiate into trabecular bone^10^. These results highlight the need for controlled, sustained proliferation of chondroprogenitors to maintain active growth plates.

Data from our group has shown that the transcription factor STAT3 regulates human chondrocyte proliferation at both fetal and adult stages^11, 12^. Chondrocytes enriched for active STAT3 are more proliferative and are also enriched for SOX9 protein, the master regulator of chondrogenesis, while evidencing negligible levels of the osteogenic transcription factor RUNX2^12^. Inhibition of either Il6ST (gp130), the obligate co-receptor for all IL-6 family cytokines, or LIFR which heterodimerizes with gp130 to facilitate LIF signaling, greatly reduced chondrocyte proliferation and STAT3 activation, respectively^11^. Inhibition of STAT3 activity in human fetal chondrocytes greatly reduced proliferation while increasing apoptosis, suggesting an important role for this molecule and the signaling mediated by gp130 in developmental chondrogenesis^11^. This is supported by data from zebrafish, in which *stat3* null fish evidence fully penetrant scoliosis, reduced proliferation and increased apoptosis^13^. Moreover, deletion of *Stat3* in mesodermal and early skeletal tissues using *T*-Cre and *Prrx1*-Cre, respectively, caused dwarfism, skeletal abnormalities and kyphosis; the authors did not assess proliferation, but did observe significantly reduced SOX9 levels in *Stat3^-/-^* chondrocytes^14^. They went on to show that stimulation with the IL-6 family cytokine oncostatin M (OSM), which can signal through a LIFR/gp130 complex and/or a OSMR/gp130 complex^15^, could increase both STAT3 activation and SOX9 levels *in vitro*. Few of these animals survived past P7, precluding the analysis of chondroprogenitors at later stages of growth plate function. Together, these studies identify an important role for STAT3 in regulating developing chondrocyte proliferation and apoptosis. However, the function of STAT3 in postnatal developing chondrocytes, the upstream regulators of this transcription factor and the potential role of STAT3 in the sexual dimorphism of growth plate closure remain to be elucidated.

Here we show that postnatal deletion of *Stat3* in chondrocytes using *Acan*-Cre^ERT2^ results in decreased proliferation, increased hypertrophy and apoptosis, stunting and growth plate fusions in the long bones and the spine. Female mice are more strongly impacted than males. We also demonstrate that chondrocyte-specific deletion of *Stat3* results in progressive dysfunction of articular cartilage, although this compartment appears to be less affected than the growth plate. Deletion of gp130, the obligate co-receptor for all IL-6 cytokines in postnatal chondrocytes mostly phenocopies deletion of *Stat3*, while deletion of *Lifr* results in a milder phenotype. Conversely, Stat3 overexpression in postnatal chondrocytes resulted in hyperproliferation and skeletal abnormalities and rescued the thin growth plates observed in *gp130* deletion mutants. Finally, we show that estradiol increases STAT3 levels and activity *in vitro* and inhibition of estrogen signaling in female mice reduces Stat3 levels and activation, suggesting a potential mechanism behind the dimorphic effects seen following *Stat3* deletion in mice. Collectively, these data define Stat3 as a regulator of postnatal chondrocyte cell proliferation, differentiation and survival and implicates IL-6 family cytokine signaling as a positive regulator of chondrogenic cell proliferation and maintenance.

## Materials and Methods

### Mouse lines and breeding

All procedures involving animals were approved by the Institutional Animal Care and Use Committee of USC. This study was compliant with all relevant ethical regulations regarding animal research. Mice with tamoxifen-regulated expression of the Cre recombinase under control of the chondrocyte-specific aggrecan (Acan) promotor (*Acan*-Cre^ERT2^) were purchased from JAX (strain 019148^16^) and used exclusively as heterozygotes (*Acan*-Cre^+/ERT2^). To delete *Stat3*, these mice were crossed with mice bearing floxed *Stat3* alleles (*Stat3^fl/fl^*), also purchased from JAX (strain 016923^17^). To overexpress Stat3, mice carrying a floxed stop cassette followed by a constitutively active *Stat3* allele in the *Rosa26* locus (Rosa26-Stat3C^fl^*^fll^*; a kind gift from Dr. Sergei Koralov of New York University School of Medicine^18^) were crossed with *Acan-*Cre^+/ERT2^ mice. To delete *Il6st* (gp130), mice bearing floxed gp130 alleles (*gp130^fl/fl^*) were purchased from the Knockout Mouse Project (KOMP) Repository of University of California, Davis (*Il6st^tm1a(KOMP)Mbp^*) and were crossed with mice expressing Flp recombinase under the control of the constitutive *Rosa26* promoter (JAX strain 009086^19^) to remove the Neo cassette. Animals negative for Flp recombinase were then crossed to *Acan-*Cre^+/ERT2^ mice. *In vivo* experiments were performed with cohoused littermate controls. *Lifr*^fl/fl^ (Lifr^tm1c(EUCOMM)Hmgu^) cryopreserved frozen sperm were purchased from MRC Harwell Institute, The Mary Lyon Centre, UK. The mice were re-derived at USC. *Col2a1*-Cre mice were purchased from JAX (strain 003554^20^). *Gli1*-Cre^+/ERT2^ mice were purchased from JAX (strain 007913^21^). All animals were on a C57BL/6 background. Genotyping was performed according to protocols provided by JAX, KOMP and MRC.

### *STAT3* knockdown and gene expression analysis

Human fetal chondrocytes from the interzone were isolated as previously described^22^ from de-identified material from anonymous donors following informed consent. Passage 0 chondrocytes were cultured *in vitro* for 2-3 days before being infected with doxycycline inducible *STAT3* shRNA or scrambled lentiviral particles (Dharmacon). One day later, doxycycline treatment commenced for a total of three days; fresh doxycycline was added daily. After three days, cells were sorted for RFP and mRNA was isolated with the RNeasy kit (Qiagen). Libraries (KAPA LTP kit, Roche) were prepared after RNA quality validation (Agilent Bioanalyzer 2100) and sequenced on a HiSeq3000 (Illumina). Raw sequencing reads were processed using Partek Flow Analysis Software. Briefly, raw unaligned reads were quality checked, trimmed and reads with a Phred quality score >20 were used for alignment. Trimmed reads were aligned to human genome version hg38 with Gencode release 29 annotations using STAR aligner (2.5.3a) with default parameters. Transcript levels were quantified to the annotation model using Partek’s optimization of the expectation-maximization algorithm using default parameters. Transcript counts were normalized using reads per kilobase million (RPKM) approach and transformed to log2(RPKM+1). Differential expression gene expression analysis was performed by the DESeq2 R package^23^.

### RNA extraction and quantitative Real-Time PCR

Total RNA was extracted using the RNeasy Mini Kit (Qiagen) and cDNA was amplified using the Maxima First Strand cDNA Synthesis Kit (Thermo Fisher). Power SYBR Green (Applied Biosystems) RT-PCR amplification and detection was performed using an Applied Biosystems Step One Plus Real-Time PCR machine. The comparative Ct method for relative quantification (2-ΔΔCt) was used to quantitate gene expression, where results were normalized to *Rpl7* (ribosomal protein L7). Primer sequences are available upon request.

### Western blot analysis

Non-treated xiphoid processes were carefully dissected from 2 month old male and female WT mice, ground and then dissolved in RIPA Lysis and Extraction Buffer (Pierce) containing protease inhibitors (Pierce) followed by sonication with a 15-second pulse at a power output of 2 using the VirSonic 100 (SP Industries Company). Protein concentrations were determined by BCA protein assay (Pierce) and boiled for 5 minutes with Laemmli Sample Buffer (Bio-Rad, Hercules, CA). Proteins were separated on acrylamide gels and analyzed by Western blot using primary antibodies: anti-STAT3 (9139), anti-pSTAT3 (9145) and anti-Histone H3 (9515; all from Cell Signaling). Histone H3 antibody was used as a loading control. Proteins were resolved with SDS-PAGE utilizing 4–15% Mini-PROTEAN TGX Precast Gels and transferred to Trans-Blot Turbo Transfer Packs with a 0.2-µm pore-size nitrocellulose membrane. The SDS-PAGE running buffer, 4–15% Mini-PROTEAN TGX Precast Gels, Trans-Blot Turbo Transfer Packs with a 0.2-µm pore-size nitrocellulose membrane were purchased from Bio-Rad. Nitrocellulose membranes were blocked in 5% nonfat milk in 0.05% (v/v) Tween 20 (Corning). Membranes were then incubated with primary antibodies overnight. After washing in PBS containing 0.05% (v/v) Tween 20 (PBST), membranes were incubated with secondary antibodies (31460 and 31430, Thermo Scientific). After washing, development was performed with the Clarity Western ECL Blotting Substrate (Bio-Rad) and images were quantified by ImageJ software.

### Immunohistochemistry (IHC)

Limb and spine tissues were dissected and fixed in 10% formalin for 24 hours. They were then decalcified with 14% EDTA, pH7.4, for 4-5 days at 4 °C, embedded in paraffin and cut at a thickness of 5 µm. Paraffin sections were deparaffinized and rehydrated by passage through xylene and 100%, 95% and 70% ethanol. Antigen Retrieval was carried out in citrate buffer (pH 6.0) for 30 min at 60 °C. Sections were blocked with 2% normal horse serum in TBS for 1 h at room temperature and incubated with primary antibody in 1% BSA at 4 °C overnight. Slides were washed 3 times in TBS containing 0.05% (v/v) Tween 20 (TBST) for 5 min each and incubated at room temperature for an hour in secondary antibody-HRP (Vector Laboratories). Antibodies were then visualized by peroxidase substrate kit DAB (Vector Laboratories). Slides were viewed using a Zeiss Axio Imager.A2 Microscope and images were taken using an Axiocam 105 color camera with Zen 2 program. Standard microscope camera settings were used. Auto-exposure was used to normalize background light levels across all images. Protein expression was quantified using ImageJ by measuring number of positively labeled cells in each section normalized to total number of analyzed cells and expressed as a percentage of positive cells. Primary antibodies included the following: aggrecan neoepitope (Novus Biologicals; NB100-74350), Runx2 (Santa Cruz Biotechnology; sc-390351), collagen X (Abcam; ab58632), SOX9 (Abcam; 26414), pStat3 (Cell Signaling; 9415), Stat3 (Cell Signaling; 9139), gp130 (Biosis; bs-1459R), Lifr (Biosis; bs-1458R) and aggrecan (Abcam; ab36861). For pStat3 IHC, EDTA antigen retrieval buffer, pH 8.0, at 90 °C for 40 min was used. Safranin O staining was performed as described^24^.

### Determination of apoptosis and proliferation

TdT-mediated dUTP nick-end labeling (TUNEL) or EdU assays were performed by using *in situ* cell death detection kit (TUNEL, Promega Corporation, G3250) or EdU Click-iT® Assay Kit obtained from Thermo Fisher (C10639), respectively, as described in the manufacturer’s protocol. EdU (Abcam) was injected intraperitoneally daily at 25 mg/kg for 4 days (*Acan*-Cre^+/ERT2^;*Stat3*^fl/fl^ mice) or 4 hours (*Acan*-Cre^+/ERT2^;Rosa26-Stat3C^fl/fl^) prior to sacrifice. Slides were counterstained with DAPI. Apoptosis and proliferation were evaluated by Image J analysis.

### MicroCT data collection and analysis

Eight mice (2 per sex per genotype; *Stat3^fl/fl^* and *Acan*-Cre^+/ERT2^;*Stat3^fl/fl^*) were microCT scanned with a GE phoenix nanotom m X-ray nanoCT scanner with the following x-ray parameters to assess axial skeleton changes: 70 kV energy, 160 µA current, 1440 projection along 360-degree rotation, 1 frame per second, averaging 2 frames and skipping 1. Three consecutive scans were performed to cover whole mouse at 0.02 mm voxel resolution. All three initial scans were reconstructed using embedded software in the scanner. Visualization of 3D morphological structures and acquisition of measurements for quantification were performed using VGSTUDIO MAX 3.3 (Volume Graphics GmbH). Landmark-based data were collected from the lumbosacral region and pelvic girdle. Eight measurements were collected for each lumbar vertebra (Supplementary Table 3).

Geometric morphometrics (GM) analysis was applied to assess phenotypic changes. A total of 34 sacral and 26 pelvic landmarks were obtained for the 3D GM analysis. The sacral landmarks utilized were: **1**. The most cranioventral point on S1 (sacral vertebra) centrum on midsagittal plane; **2**. The most craniodorsal point on S1 centrum on midsagittal plane; **3**. The right ventrolateral point on S1 centrum outline cranially, where the ascendant part of ala starts at pedicle; **4**. The left ventrolateral point on S1 centrum outline cranially, where the ascendant part of ala starts at pedicle; **5**. The right dorsolateral point on S1 centrum outline cranially, where the ascendant part of ala starts at pedicle; **6**. The left dorsolateral point on S1 centrum outline cranially, where the ascendant part of ala starts at pedicle; **7**. The most cranial middle point on the right ala; **8**. The most cranial middle point on the left ala; **9**. The most cranioventral point on the right ala; **10**. The most cranioventral point on the left ala; **11**. The most craniodorsal point on the right ala; **12**. The most craniodorsal point on the left ala; **13**. The most caudal (deepest) point on the concave rim between the right ala and articular process; **14**. The most caudal (deepest) point on the concave rim between the left ala and articular process; **15**. The most craniomedial point on the right articular process; **16**. The most craniomedial point on the left articular process; **17**. The most craniodorsal point on the right articular process; **18**. The most craniodorsal point on the left articular process; **19**. The most cranial point on the dorsomedial S1 sacral canal wall; **20**. The most cranioventral point on S2 centrum on midsagittal plane; **21**. The most cranioventral point on S3 centrum on midsagittal plane; **22**. The most cranioventral point on S4 centrum on midsagittal plane; **23**. The most caudoventral point on S4 centrum on midsagittal plane; **24**. The most craniodorsal point on S1 neural spine; **25**. The most craniodorsal point on S2 neural spine; **26**. The most caudodorsal point on S2 neural spine; **27**. The most craniodorsal point on S3 neural spine; **28**. The most caudodorsal point on S3 neural spine; **29**. The most craniodorsal point on S4 neural spine; **30**. The most caudodorsal point on S4 neural spine; **31**. The most caudodorsal point on S4 centrum on midsagittal plane; **32**. The right caudolateral point on S4 centrum on midcoronal plane; **33**. The left caudolateral point on S4 centrum on midcoronal plane; **34**. The most caudal point on the dorsomedial S4 sacral canal wall.

Pelvic landmarks utilized were: **1**. The central point of iliac spine rugosity, site attachment for m. rectus femori; **2**. The caudodorsal point on the auricular surface (sacroiliac articular surface); 3. The ventral point on the lower ilium at the same transversal cross section as landmark 2, the transversal section corresponds to the longitudinal axis of pelvis; **4**. The lateral point on the lower ilium at the same transversal cross section as landmark 2, the transversal section corresponds to the longitudinal axis of pelvis; **5**. The medial point on the lower ilium at the same transversal cross section as landmark 2, the transversal section corresponds to the longitudinal axis of pelvis; **6**. The deepest point on the ischium where its process meets acetabulum; **7**. The ischial spine; **8**. The caudodorsal point on ischial tuberosity; **9**. The caudolateral point on the ischiopubic ramus; **10**. The caudomedial point on the ramus of the ischium; **11**. The most caudal point on the pubic symphysis; **12**. The most cranial point on the pubic symphysis; **13**. The ventral point of the ilio-pubic eminence; **14**. The deepest point on the angulation between ventral point of ilio-public eminence and the longitudinal axis of ilium; **15**. The most caudoventral point on the auricular surface; **16**. The most ventral point on the iliac crest; **17**. The most lateral point on the iliac crest; **18**. The most dorsal point on the iliac crest; **19**. The most cranial point on the rim of the obturator foramen; **20**. The most ventral point on the rim of the obturator foramen; **21**. The most dorsal point on the rim of the obturator foramen; **22**. The cranial point on the rim of articular surface of acetabulum, which is defined as a parallel projection from landmark 1 along the major axis of pelvis; **23**. The caudal point on the rim of articular surface of acetabulum, directly across from landmark 8 along the long axis of the ischium; **24**. The caudoventral point of the lunate surface of the of acetabulum; **25**. The deepest center point of the acetabulum. **26**. The craniodorsal point of the articular surface of the acetabulum, which is defined as the extension of the line connecting landmark 24 and 25.

Amira 2019.4 (FEI, Visualization Sciences Group) software was used to acquire the x, y and z coordinates for each landmark with accuracy of landmark size 0.1-0.2 mm. The visualization pattern of shape variance was performed using the interactive “MORPHOTOOLS” software^25–27^. The 3D coordinates of each landmarks were subjected to a generalized Procrustes analysis (GPA)^28, 29^, which included translating, rotating, reflecting, and scaling them to a best least-squares fitting (GLS)^30, 31^. Variation among landmark configurations and shape were explored via principal component analysis (PCA)^32^. Thin-plate spline analysis was used to warp surfaces on the basis of reference configuration and visualize the shape difference along the principal axis.

Three mice per sex per genotype (*Stat3^fl/fl^* and *Acan*-Cre^+/ERT2^;*Stat3^fl/fl^)* were microCT scanned for epiphyseal trabecular bone analysis. Image data were acquired as above and focused on the knee joint to obtain 10 µm pixel size resolution images in order to be able to calculate trabecular properties on the proximal epiphyseal segment of tibia. The X-ray parameters used for these 10 µm pixel size resolution images included: 70kV energy, 120uA current, and a voxel resolution of 10 µm. Each specimen rotated 360 degrees about a vertical axis in front of the detector resulted in 1,440 projections acquired at one frame per second. The 1,440 projections were reconstructed into 16-bit unsigned images saved as .dicom formatted files using proprietary software of Nanotom GE phoenix (i.e., datosx 2 rec). The dicom files were used as the basis for data analyses. Visualization of 3D morphological structures and acquisition of measurements for quantification were performed using VGSTUDIO MAX 3.4 64 bit (Volume Graphics GmbH).

After acquisitions of the initial image data on the knee joint, a subset of region of interest (ROI) in the proximal tibial epiphysis was created; cortical bone was removed from epiphysis up to the most distal limit of medial and lateral epicondylar transversal line. From this ROI trabecular BV/TV was calculated.

For microCT analysis of *Col2a1*-Cre;*Stat3*^fl/fl^ and control animals (3 per sex per genotytpe), a Skyscan1172 (Bruker MicroCT, Kontich, Belgium) machine using CTAn (v.1.13.2.1) and CTVol (v.2.3.2) software was employed. The microradiography unit was set to an energy level of 55 kVp, intensity of 181 µA, and 900 projections; specimens were scanned at an 8 µm voxel resolution. A three-dimensional reconstruction was generated with NRecon software (Bruker MicroCT) from the set of scans. Femurs were measured directly from these reconstructions.

### Tissue collection and digestion

Mouse knee joints were cut at the femur and the tibia and crushed lightly with a mortar and pestle and placed in digestion media (DMEM/F12 (Corning) with 10% Fetal Bovine Serum (FBS; Corning), 1% penicillin/streptomycin/amphotericin B solution (P/S/A; Corning), 1 mg/mL dispase (Gibco), 1 mg/mL type 2 collagenase (Worthington), 10 µg/mL gentamycin (Teknova) and 100 μg/mL primocin (Invivogen)) in an Erlenmeyer flask with a spin bar in 4 °C overnight followed by 37 °C for 4 - 6 hours. Cells were washed with DPBS (Corning) after digestion and strained through a 70 μm filter (Fisher) before FACS.

### FACS

All FACS was performed on a BD FACSAria IIIu cell sorter. Mouse knee cells were washed twice in 1 - 2% FBS and stained with DAPI for viability. Populations of interest based on DAPI negativity expression and lack of lineage markers (APC-Cy7; CD31, Ter119, CD45 at 1 μL/10^6^ cells; Biolegend) expression were directly sorted into DMEM/F12 containing 10% FBS with 1% P/S/A. Flow cytometry data was analyzed using FlowJo software.

### Single-cell sequencing using 10X Genomics

Single cell samples were prepared using Chromium Next GEM Single Cell 3’ Reagent Kits v3.1 and chip Kit (10X Genomics) according to the manufacturer’s protocol. Briefly samples were FACS sorted using DAPI to select live cells followed by resuspension in 0.04% BSA-PBS. Nearly 1200 cells/µl were added to each well of the chip with a target cell recovery estimate of 10,000 cells. Thereafter Gel bead-in Emulsions (GEMs) were generated using GemCode Single-Cell Instrument. GEMs were reverse transcribed, droplets were broken and single stranded cDNA was isolated. cDNAs were cleaned up with DynaBeads and amplified. Finally, cDNAs were ligated with adapters, post-ligation products were amplified, cleaned-up with SPRIselect and purified libraries were sequenced on Novaseq at the UCLA Technology Center for Genomics and Bioinformatics.

### 10X sequencing data analysis

Raw sequencing reads were processed using Partek Flow Analysis Software (build version 10.0.21.0602). Briefly, raw reads were checked for their quality, trimmed and reads with an average base quality score per position >30 were considered for alignment. Trimmed reads were aligned to the mouse genome version mm10 using STAR-2.7.3a with default parameters. Reads with alignment percentage >75% were de-duplicated based on their unique molecular identifiers (UMIs). Reads mapping to the same chromosomal location with duplicate UMIs were removed. Thereafter ‘Knee’ plot was constructed using the cumulative fraction of reads/UMIs for all barcodes. Barcodes below the cut-off defined by the location of the knee were assigned as true cell barcodes and quantified. Further noise filtration was done by removing cells having >5% mitochondrial counts and total read counts >24,000. Ambient mRNAs were removed using the R package SoupX^33^. Genes not expressed in any cell were also removed as an additional clean-up step. Cleaned-up reads were normalized using counts per million (CPM) method followed by log-transformation generating count matrices for each sample. Samples were batch-corrected on the basis of expressed genes and mitochondrial reads percent. Count matrices were used to visualize and explore the samples in further details by generating tSNE plots, a non-linear dimensional reduction technique. This algorithm learns the underlying manifold of the data and places similar cells together in a low-dimensional space. Dot plots were generated in R (v 4.0.3) using ggplot2 (v3.3.3).

### Gene set enrichment analysis (GSEA)

Single cell sequencing data was analyzed using the GSEA-P desktop application^34^ available at www.broadinstitute.org. The specific parameters used for the analyses are: number of permutations (1000), permutation type (gene set), enrichment statistic (weighted), metric for ranking genes (signal to noise). Enrichment score was calculated by walking down the rank ordered gene list from top to last ranked gene, decreasing a running-sum statistic when a gene is encountered that is not part of a gene set, and increasing the score when a gene is encountered that is in the gene set. Hypertrophic gene set included genes *Ptgis, Snai1, Cd200, Plin2, Foxa2, Ugp2, Adk, Lxn, Enpp2, Mef2c, Col10a1, Sp7, Alpl, Ihh* and *Col1a1*. Proliferative gene set included genes *Itga4, Cdk1, Dcx, Sox5, Fn1, Cdk2, Sox6, Ptges3, Foxn2, Sdc4, Ncam1, Rps25, Lmna, Bmpr1b* and *Prg4*.

### Statistical analysis

Numbers of repeats for each experiment are indicated in the figure legends. Pooled data are represented as mean ± SEM unless otherwise indicated. Statistical analysis was performed using 2-tailed Student’s t test to compare 2 groups unless noted, in which case ANOVA was used to compare 3 or more groups. p values less than 0.05 were considered to be significant.

## Results

### Stat3 promotes proliferation and survival in postnatal growth plate chondrocytes

During development, chondrocytes form a highly proliferative blastema to supply rapidly growing limbs^22^. We have recently published a survey of the human chondrocyte transcriptome throughout ontogeny, spanning early embryonic development through adult stages^12^. Interrogation of this longitudinal data set for IL-6 family cytokines and receptors, known regulators of STAT3 activity, demonstrated consistent expression of gp130 and stage-specific expression of *LIF* to fetal and adolescent stages, during which growth plates are active in humans (Supp. Fig. 1). To understand the function of STAT3 during these stages, we knocked down *STAT3* gene expression with lentiviral shRNA in human fetal chondrocytes isolated from the interzone at 15-17 weeks of development and compared their transcriptome to control cells (Fig. 1a). Gene ontology (GO) analysis^35, 36^ of the genes significantly enriched (>1.5 fold, p<0.05) in control vs. *STAT3*-depleted cells yielded categories related to matrix anabolism and secretion as well as proliferation, confirming our earlier results^11^ and further suggesting a relationship between STAT3 activity and chondrocyte proliferation and matrix protein production. Genes downregulated upon *STAT3* knockdown included *ACAN*, *COL2A1*, *MATN3* and *MATN4* (Supplementary Table 1). To assess the function of STAT3 *in vivo*, we crossed *Stat3*^fl/fl^ mice^37^ with the *Acan-*Cre^ERT2^ strain^16^, enabling tamoxifen-inducible deletion of *Stat3* in chondrocytes with a single allele of *Acan-*Cre^ERT2^. Pups were administered tamoxifen at postnatal days 2 and 3 (P2 and P3) by gavage and then skeletal phenotypes were assessed at stages corresponding to sexual immaturity and maturity, skeletal maturity and early aging (1, 3, 6 and 9 months of age, respectively; Fig. 1b,c). Compared to *Stat3*^fl/fl^ littermate controls, *Stat3* mutants survived at equivalent rates but exhibited significant decreases in body weight and were proportionally smaller at all ages examined. Notably, females were more strongly affected then males at 3 and 9 months and evidenced a non-significant trend at 6 months as well (Fig. 1d); levels of *Acan* mRNA and protein were not different between females and males at the time of induction (Supp. Fig. 2). To assess if there were sex-specific differences in the levels of Stat3 protein, we stained the growth plates of wild type female and male mice for Stat3 using immunohistochemistry (IHC); these data demonstrated that not only was there more Stat3 in female growth plates, but in both sexes Stat3 was mostly localized to the resting and proliferative zones of the growth plate (Fig. 1e). We then isolated sternal chondrocytes from the xiphoid process of both female and male wild type mice and quantified protein by Western blot. These data (Fig. 1f) demonstrated a clear enrichment for both total and active (phosphorylated; p) Stat3 in female mice, which may contribute to the observed dimorphism in reliance upon *Stat3* for growth.

**Figure 1:**
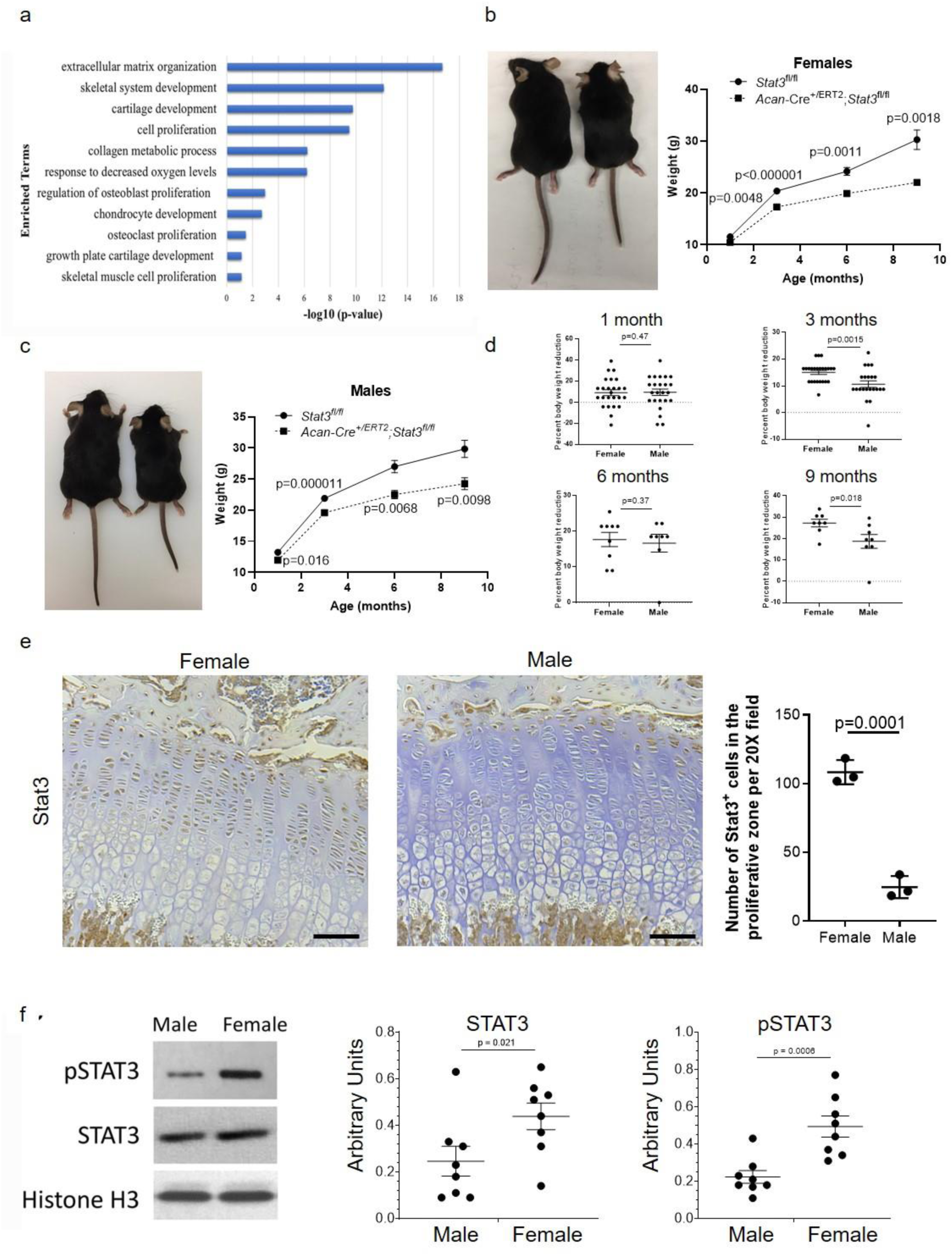
STAT3 regulates anabolic genes in human fetal chondrocytes and body size in mice. (a) Altered gene expression in human fetal chondrocytes after knockdown of STAT3 by shRNA *in vitro* as determined by bulk RNA-Seq and GO analysis; n=3. (b, c) Postnatal deletion of *Stat3* in chondrocytes via *Acan*-Cre^+/ERT2^ *in vivo* decreases body weight and size of female (b) and male(c) mice; n≥7 for each sex and genotype per timepoint. Images of 6 month old mice are shown. (d) Reductions in body weight in *Stat3* mutants compared to WT littermates were greater in female than in male mice. (e) Stat3 protein localized to the resting and proliferative zones of the growth plate at 1 month in WT mice. (f) Stat3 and pStat3 levels were higher in WT female mice as compared to WT male mice (n=8). Scale bars = 50 µm.

Based on the size differences between *Acan-*Cre^+/ERT2^;*Stat3*^fl/fl^ mice and controls as early as 1 month, we measured the width of each zone in the growth plate in control and *Stat3* deletion mutants at this age (Fig. 2a,b); given the potential for insertion of Cre^ERT2^ into the *Acan* locus to disrupt skeletal development, we also verified that mice bearing a single allele of *Acan-*Cre^ERT2^ were not affected as has been previously published^38, 39^. These data revealed no differences between *Acan*-Cre^+/ERT2^;*Stat3*^fl/fl^ and control animals in the resting zone where chondroprogenitors localize^9^; however, the length of the proliferative zone was significantly reduced in *Stat3* deleted animals, concurrent with an increase in the pre-hypertrophic and hypertrophic zones (Fig. 2a,b). To validate the independence of the observed phenotype from the Cre line used to delete *Stat3,* we also generated both *Col2a1*-Cre;*Stat3*^fl/fl^ as well as *Gli1*-Cre^+/ERT2^;*Stat3* ^fl/fl^ mice. In the former, Cre is expressed constitutively in chondrocytes, resulting in *Stat3* deletion during mid-embryogenesis. At 3 months, both *Col2a1*-Cre;*Stat3*^fl/fl^ female and male mice were visibly smaller than *Stat3*^fl/fl^ control animals, and microCT measurements of femoral length demonstrated significant shortening of femurs in *Stat3* deleted animals (Supp. Fig. 3a); there was a non-significant trend for females to be more impacted than males (Supp. Fig. 3b). Growth plates were also impacted, with significantly reduced widths and bony bridges evident (Supp. Fig. 3c,d). In *Gli1*-Cre^+/ERT2^;*Stat3* ^fl/fl^ animals induced with tamoxifen at P2/P3 analyzed at 1 month of age, mice were visibly smaller and growth plates were significantly thinner in both females and males as compared to controls (Fig. 2d-f).

**Figure 2:**
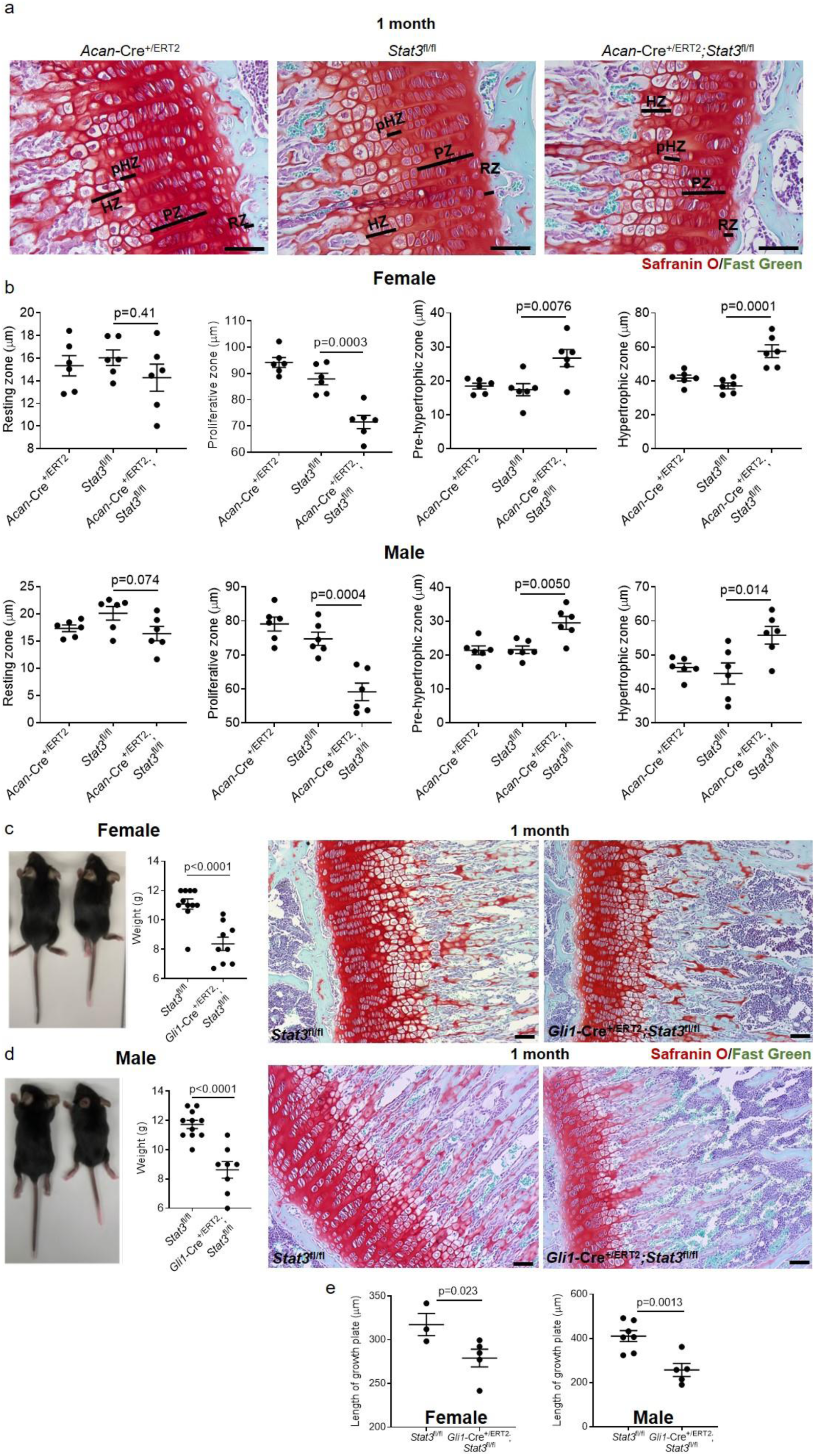
Postnatal loss of *Stat3* in chondrocytes decreases the thickness of the growth plate at 1 month. (a,b) Following deletion of *Stat3* at P2/P3 via *Acan*-Cre^+/ERT2^, the proliferative zone of the proximal tibial growth plate was significantly reduced, while the pre-hypertrophic and hypertrophic zones were increased, in both female and male mice; the resting zone was not affected. *Acan*-Cre^+/ERT2^ animals treated with tamoxifen did not show any reduction in thickness of growth plate zones. n = 6 animals per genotype, p values were calculated with one-way ANOVA test. (c) Deletion of *Stat3* at P2/P3 in *Gli1*^+^ cells resulted in reduced body weight and significantly smaller growth plates in both female (c,e) and male (d,e) mice. Scale bars = 50 µm.

To assess the cellular basis for the observed growth retardation and disruptions in the zonal architecture of the growth plate, we measured the thickness of the proximal tibial growth plates at multiple stages following early postnatal deletion of *Stat3*. At one month of age, both female and male mice lacking *Stat3* demonstrated reductions in total growth plate thickness; these deficits persisted with advancing age. Beginning at 3 months we observed growth plate fusions in *Stat3* mutants of both sexes (Fig. 3a,b), which were significantly more frequent in females at 3 months (Supp. Fig. 3e). These fusions were more frequent in older animals, suggesting a progressive loss of growth plate chondrocytes and potential replacement by bone. To validate this, we conducted microCT analysis of *Acan*-Cre^+/ERT2^;*Stat3*^fl/fl^ and *Stat3*^fl/fl^ control animals at 6 months following postnatal deletion at P2/P3; this analysis revealed significantly greater bone density in the proximal tibial epiphysis of *Stat3* deleted females and males compared to controls (Supp. Fig. 4). To better assess the cellular basis underlying thinner growth plates and the reduced size of the proliferative zone observed at 1 month, we injected EdU into 3 month old animals for 4 consecutive days before analysis (Fig. 3d); these data showed significantly fewer proliferating cells in the growth plates of *Acan*-Cre^+/ERT2^;*Stat3*^fl/fl^ animals as compared to *Stat3*^fl/fl^ controls. Finally, based on the acknowledged function of Stat3 as an anti-apoptotic factor^11, 17^ we assessed the rate of cell death in growth plates of *Stat3* deleted and control animals at 6 months following tamoxifen administration at P2/P3. *Stat3* null chondrocytes evidenced significantly more apoptosis (Fig. 3e). Together, these data support a role for Stat3 in promoting chondrocyte proliferation and survival to maintain growth rates of the long bones, and also suggest a role for Stat3 in inhibiting epiphyseal bone formation in the distal region of the growth plate.

**Figure 3:**
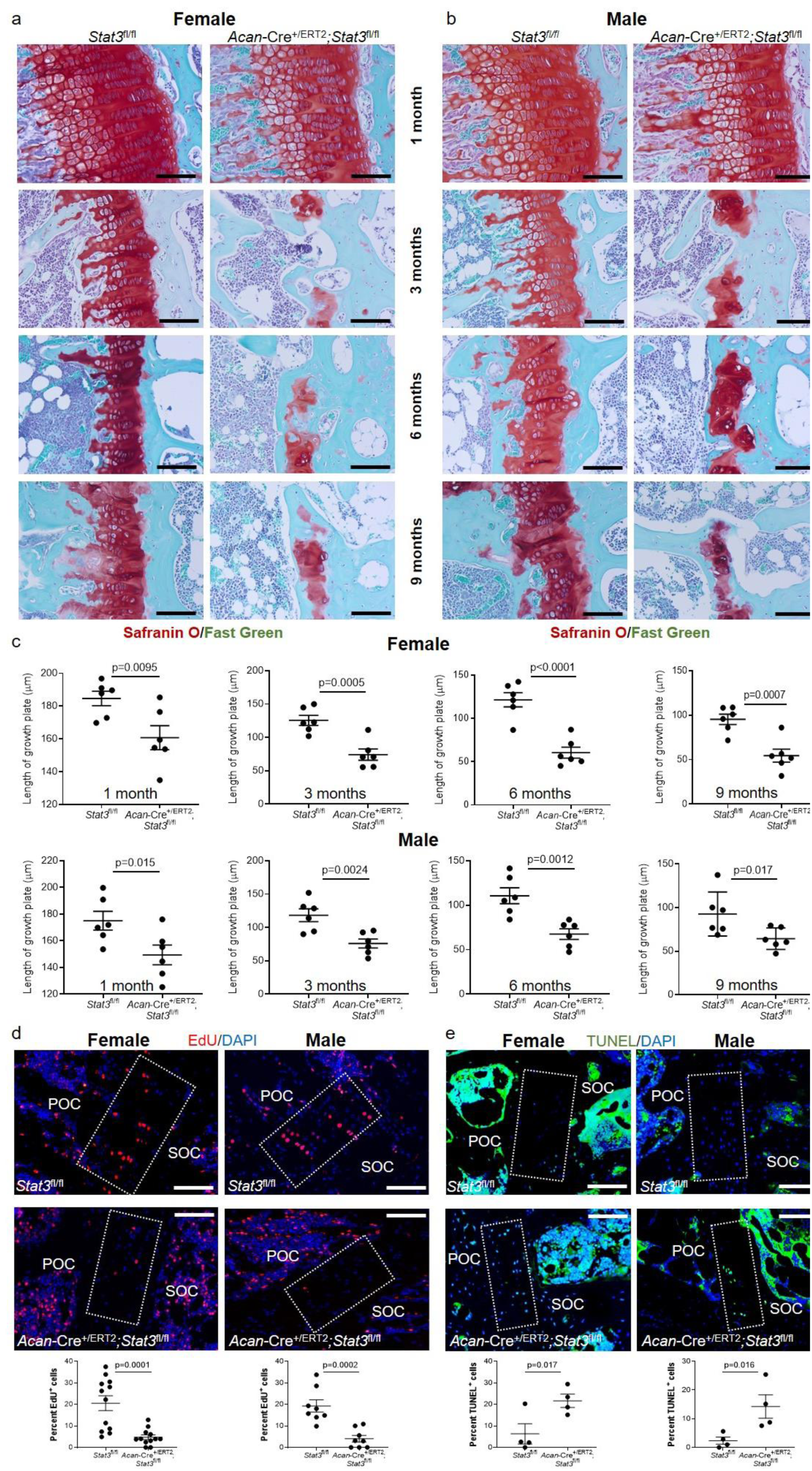
Postnatal loss of *Stat3* in chondrocytes decreases proliferation and increases apoptosis. Proximal tibial growth plates were greatly reduced in thickness following deletion of Stat3 at P2/P3 in (a,c) female and (b,c) male mice. Beginning at 3 months, growth plate fusions were observed with replacement by trabecular bone. Scale bars equal 50 µm. (d) EdU was injected intraperitoneally for 4 days before analysis of mice at 3 months of age. EdU^+^ cells in the proximal tibial growth plate were scored. (e) Apoptosis of chondrocytes in the proximal tibial growth plate (3 month old) was detected by TUNEL. Boxed regions indicate the growth plate. For each group, n=6. Scale bars = 50 µm. POC = primary ossification center; SOC = secondary ossification center.

### Stat3 is required for vertebral end plate integrity

In the spine, chondroprogenitors located in the vertebral end plates produce chondrocytes that form growth plates^40, 41^; this mechanism is very similar to that found in long bones. Based on the shorter stature of *Stat3* deleted mice (Fig. 1), we examined vertebral elements in *Acan*-Cre^+/ERT2^;*Stat3^fl/fl^* and control *Stat3^fl/fl^* mice at several ages following induction with tamoxifen at P2/P3. *Acan-*Cre^ERT2^ targets the endplates, annulus fibrosis, and nucleus pulposus^16^. Similar to the tibial growth plate, Stat3 protein activation was localized to the proliferative zone and was enriched in female vs. male mice at 1 month (Supp. Fig. 5). Moreover, the height of the vertebral bodies was significantly shorter in *Stat3* deleted mice as early as one month of age (Fig. 4a-d). Control animals had organized vertebral structures at all stages examined between 1 and 9 months. Cartilaginous endplates had clear, continuous borders and the formation of annular lamellae was observed. The nucleus pulposus had a rich proteoglycan matrix with sparse notochordal cells. In contrast in *Stat3* deleted mice, starting at 3 months we observed a loss in continuity of both cranial and caudal endplates (Fig. 4c,d). Structural degradation of the annulus was evidenced by a loss of collagen-rich lamellae. Within the nucleus pulposus, degradation was characterized by the loss of proteoglycan matrix, an appearance of large void spaces and loss of apparent nucleus and notochordal cell morphology. The progressive loss of endplate integrity in *Stat3* mutants, followed by alterations in the nucleus pulposus, were strikingly similar to those observed in mice with postnatal deletion of *Sox9* via *Acan-*Cre^ERT2^ ^42^.

**Figure 4:**
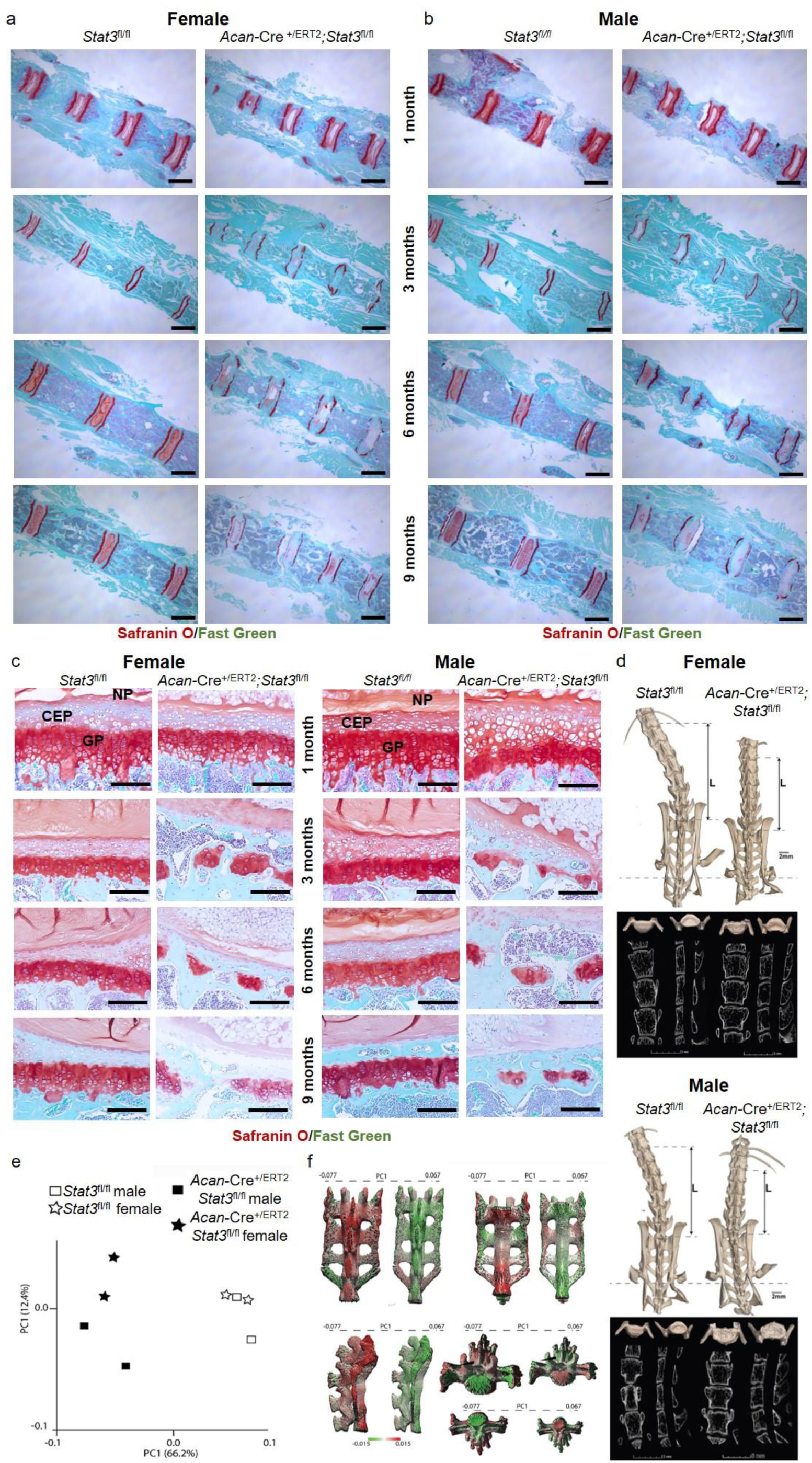
Postnatal deletion of *Stat3* affects vertebral growth plates and skeletal morphology. (a) Coronal sections of caudal vertebrae of *Acan*-Cre^+/ERT2^;*Stat3*^fl/fl^ and control *Stat3*^fl/fl^ mice demonstrate shorter vertebral bodies in both female and (b) male mice, coupled with (c) loss of vertebral growth plates. Scale bars = 1 mm (a,b) and 50 µm (c); n=6. AF = annulus fibrosis, CEP = cartilaginous end plate and NP = nucleus pulposus. (d) MicroCT and X-ray imaging of the lumbar and sacral vertebrae of control female (top panel, left), *Stat3* deleted female (top panel, right), control male (lower panel, left) and *Stat3* deleted male (lower panel, right) mice at 6 months of age following postnatal deletion at P2/P3. Note the shorter, wider vertebral bodies in *Stat3* deleted animals, resulting in overall shorter lumbar length (L). See Supplementary Table 2 for quantification. n=2. (e) Principal component analysis (PCA) of sacra from *Stat3* deleted and control mice based on distances between 34 anatomical landmarks (Supp. Fig. 5). Animals clearly segregate based on genotype. (f) Deviations along PC1 are shown on 3D surface maps of control (left) and *Stat3* deleted (right) male sacral regions. Maps are colored based on degree of deformation with respect to PC1 values, with red representing features more prominent in control and green more prominent in *Stat3* deleted animals, respectively.

We conducted microCT imaging to characterize the extent of the axial skeletal phenotype in *Stat3* mutant mice. *Stat3* deleted animals showed a reduction in the vertebral height in both lumbar and sacral regions, as well as changes in endplate structure (Fig. 4d). This was consistent in both genders. Both anterior and posterior vertebral heights from L1 to L6 were reduced in the *Stat3* mutants compared to controls (Supp. Tables 2 and 3), ranging from 19-37% reduction. Mice lacking *Stat3* had a larger endplate surface area for both cranial and caudal endplates compared to control mice (Supp. Table 2), ranging from 9-31% larger. Intervertebral disc anteroposterior and transverse length did not differ between *Stat3* deleted and control animals. We then conducted geometric morphometric mapping of the sacrum and pelvis to gain additional understanding of how loss of *Stat3* in chondrocytes impacted overall skeletal form. For the sacrum, 34 anatomical landmarks were defined based on current best practices^43–45^ and their exact 3D coordinates defined (see *Materials and Methods*, Supp. Fig. 6). Variation among landmark configurations and shape were explored via principal component analysis^46^ (PCA; Fig. 4e). Mice were clearly segregated along PC1 based on their genotype, confirming widespread changes in skeletal morphology following early postnatal deletion of *Stat3.* To visualize these differences, thin-plate spline analysis^47^ was used to project aberrations from normal on 3D renderings of control and *Stat3* deleted male sacra (Fig. 4f). A similar analysis of pelvic structure confirmed broad and consistent skeletal malformations in mutant animals (Supp. Fig. 7). These “heat map” images provide a quantitative visual representation of how specific skeletal structures change following the loss of *Stat3*, reinforcing the role *of Stat3* in contributing to regulated chondrocyte proliferation and survival in growth plates throughout the body.

### Stat3 is required for maintenance of articular cartilage

In postnatal mouse articular cartilage, chondrocyte proliferation rates are in the low single digit range^48^; cartilage expands via increases in cell volume and matrix production rather than cell division. Our *in vitro* data from human chondrocytes suggest STAT3 increases expression of ECM genes (Fig. 1a). Thus, we hypothesized Stat3 may function in articular cartilage maintenance by supporting matrix protein production and stability. Analysis of articular cartilage in *Acan*-*Cre^+/ERT2^;Stat3*^fl/fl^ mice revealed a progressive loss of proteoglycans at the articular surface compared to control mice (Fig. 5a,b). This phenotype was exacerbated in female mice, in agreement with the results from the growth plate analysis. Loss of Safranin O was clearly evident beginning at 3 months of age in females, compared to 6 months in males. Almost all Safranin O^+^ matrix was lost in *Stat3* deleted mice by 6 months of age in females vs. 9 months for males. Concomitant with accelerated loss of proteoglycans, *Stat3* deletion resulted in increased detection of aggrecan neoepitope, an antigen present when the proteoglycan aggrecan is cleaved and degraded, at 6 months of age (Fig. 5c,d). In addition, substantial apoptosis was detected in both sexes at 9 months (Supp. Fig. 8), We also detected increased expression of markers of hypertrophy in the absence of *Stat3* including Runx2 and collagen X, indicating a mild degenerative process present in the joint^49^, coupled with decreased expression of Sox9 (Supp. Fig. 8). However, no obvious signs of articular cartilage surface fibrillations, osteophytes or joint inflammation was observed at any of the investigated stages. Together, these data describe a role for Stat3 in articular chondrocytes in maintaining ECM homeostasis and preventing adoption of a hypertrophic, arthritic-prone state.

**Figure 5:**
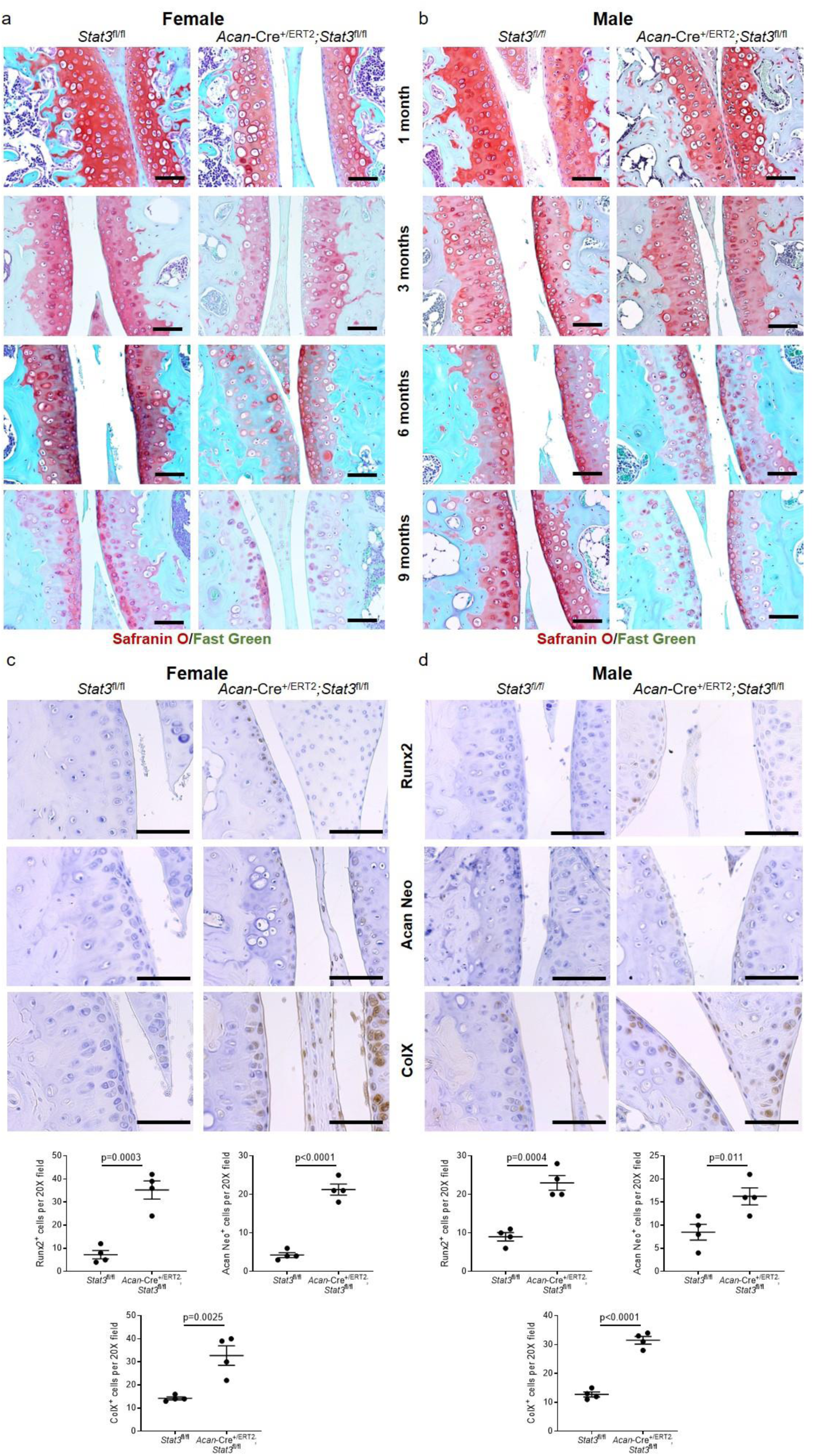
Postnatal *Stat3* deletion results in subtle changes in articular cartilage of the knee joint. (a) Administration of tamoxifen at P2/P3 to *Acan*-Cre^+/ERT2^;*Stat3*^fl/fl^ and control *Stat3*^fl/fl^ female and (b) male mice elicited reduced proteoglycan staining in articular articular cartilage by 3 months of age in female mice and by 6 months of age in males. (c) Histological staining and quantitative assessment of pre-hypertrophic (Runx2, collagen 10, COLX) and degenerative changes (aggrecan neoepitope, ACAN NEO) in the articular cartilage of knee joints of female and (d) male *Stat3* deleted and control mice at 6 months. In all panels, representative images are shown and scale bars = 50 µm. n=4-6.

### Stat3 inhibits hypertrophy and maintains a proliferative state in Sox9^+^ chondrocytes

Chondrogenesis is dependent on transcription factors of the Sox family. Sox9 is essential for chondrogenesis^50^, while Sox5 and Sox6 act downstream of Sox9 to promote proliferation and inhibit hypertrophy of chondrocytes^51^. In order to better assess the molecular consequences of postnatal *Stat3* deletion, we sorted cells negative for blood and endothelial markers from *Acan*-Cre^+/ERT2^;*Stat3*^fl/fl^ and control *Stat3*^fl/fl^ knee joints and performed scRNA-seq; we first verified the dissection strategy would maximize capture of chondrocytes by examining *Sox9*-GFP mice^52^ (Fig. 6a). Knee joints from 2-3 female mice of each genotype were digested and pooled, and live, lineage-negative cells were sorted (Fig. 6b) and analyzed at the single cell level. We focused our analysis on chondrocytes defined by expression of *Sox9* and *Col2a1*; unbiased clustering of chondrocytes expressing both genes from *Acan*-Cre^+/ERT2^;*Stat3*^fl/fl^ and control *Stat3*^fl/fl^ animals via a tSNE plot suggested some potential differences between the genotypes (Fig. 6c). Recently, Li et al. analyzed growth plate zone-specific gene regulation at the single cell level at postnatal day 7^53^; they identified genes whose expression was enriched in the proliferative and pre-hypertrophic/hypertrophic zones. Based on our data showing increases in the hypertrophic zone and bone density in the proximal epiphysis of *Acan*-Cre^+/ERT2^;*Stat3*^fl/fl^ mice compared to controls, we assessed expression levels of genes identified by Li et al., as well as known regulators of each zone, in the scRNA-seq data generated here. This analysis demonstrated highly significant enrichment of genes associated with chondrocyte maturation and hypertrophy in the absence of *Stat3* (Fig. 6d,f), while genes such as *Sox5/6* and immature chondrocytes were significantly enriched in control *Stat3*^fl/fl^ chondrocytes (Fig. 6e,f). Together, these data confirm at the transcriptional level that *Stat3* is required to suppress chondrocyte maturation and maintain a more progenitor-like state.

**Figure 6:**
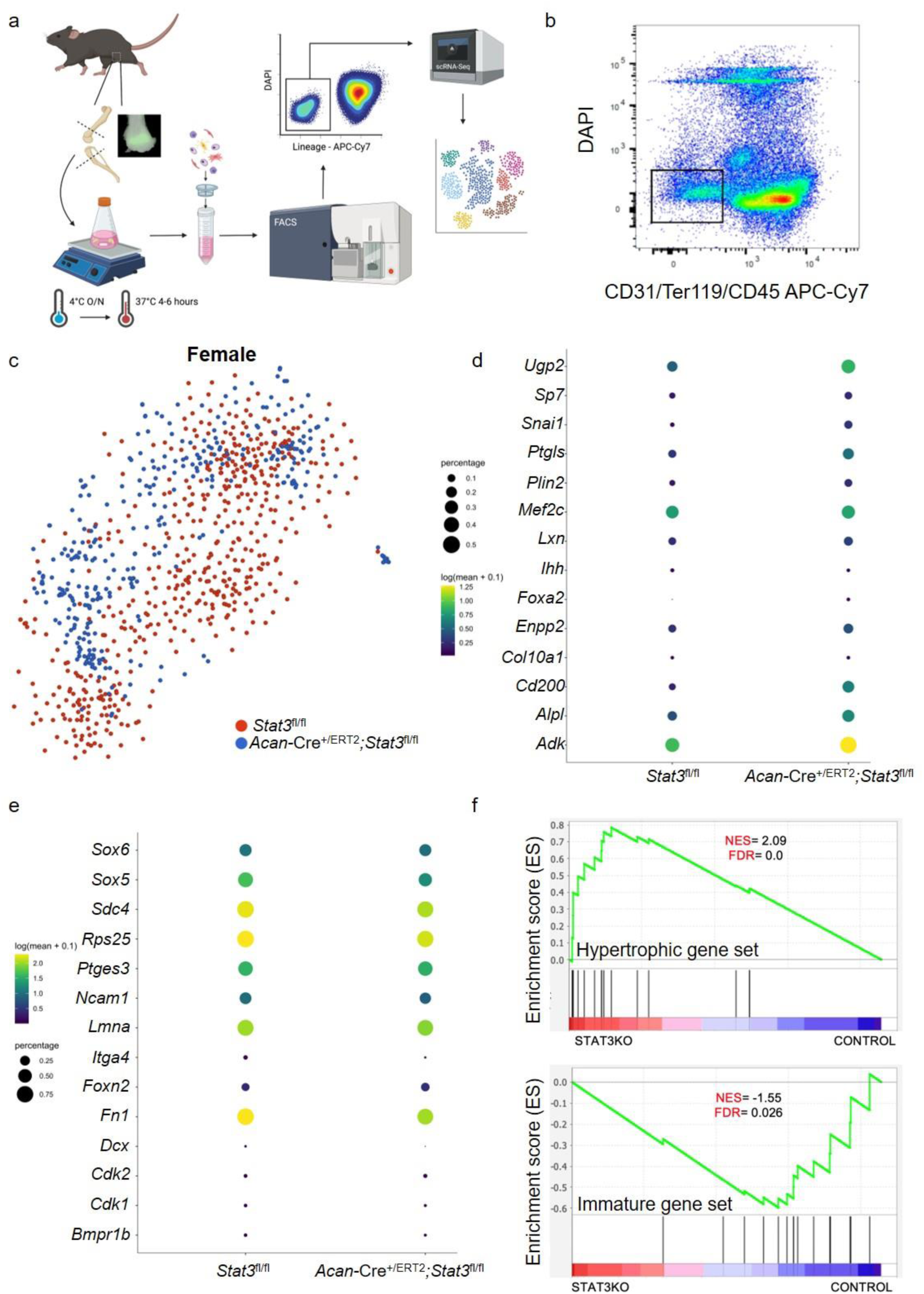
scRNA-sequencng reveals loss of *Stat3* in growth plate chondrocytes promotes premature differentiation. (a) Schematic of isolation, sorting and single cell sequencing of chondrocytes from 1 month mouse knee joints. Epifluorescent analysis of *Sox9*-GFP femurs confirmed the dissection strategy. (b) Live cells negative for markers of endothelium (CD31), red blood cells (Ter119) and white blood cells (CD45) were sorted (black box) from dissociated knee joints pooled from 2-3 female control *Stat3*^fl/fl^ or *Acan*-Cre^+/Cre^;*Stat3*^fl/fl^ mice and subjected to scRNA-sequencing. (c) tSNE plot of single cells selected for expression of *Sox9* and *Col2a1* and colored by genotype of origin. Expression pattern of genes associated with growth plate chondrocyte maturation (d) as well as immature, proliferating chondrocytes (e) are shown. (f) Gene set enrichment analysis (GSEA) plots for chondrogenic maturation (top) and immature, proliferative chondrogenic (bottom) gene sets in *Stat3*^fl/fl^ (control) and *Acan*-Cre^+/Cre^;*Stat3*^fl/fl^ (Stat3KO) female mice . The ”barcode graph” (lower portion) of each plot shows the target genes rank-ordered (left to right) by their differential expression in Stat3KO mice as compared to control mice (indicated by Enrichment Score on the y-axis). Signal-to-noise ratio metric was used for ranking of gene expression patterns in cells. NES = normalized enrichment score; FDR = false discovery rate.

### Stat3 acts downstream of gp130 to maintain output from immature chondrocytes

Many receptor tyrosine kinases, including EGFR^54^, PDGFR^55^ and GHR^56^ among others, can activate STAT3 through various signaling cascades. Based on our previous *in vitro* studies in articular chondrocytes^11^, we hypothesized that gp130, an obligate receptor for IL-6 family cytokines, may function upstream of Stat3 activation in immature chondrocytes. We found that RCGD 423, a small molecule modulator of gp130 that increases activation of Stat3, increased proliferation of articular chondrocytes^11^ without activation of extracellular matrix catabolism. Analysis of gp130 expression in the growth plate of wild type mice at 1 month demonstrated enrichment in the resting and proliferative zones, similar to the expression pattern of Stat3 (Supp. Fig. 9a). Accordingly, we crossed *Acan*-Cre^+/ERT2^ mice to *gp130*^fl/fl^ mice^57^ and administered tamoxifen at P2/P3 to ablate *gp130* postnatally in chondrocytes. At all ages examined, there was significantly reduced growth plate thickness in both female and male *gp130* deleted mice compared to *gp130*^fl/fl^ controls, including frequent disruptions in growth plate integrity by 3 months (Fig. 7a-c); as was observed in *Stat3* deleted animals, growth plate fusions were significantly more frequent in females at 3 months (Supp. Fig. 9b). Quantification of growth plate zones at 1 month yielded similar results to deletion of *Stat3* in chondrocytes, demonstrating a highly significant decrease in the thickness of the proliferative zone and increases in the more mature pre-hypertrophic and hypertrophic zones in *gp130* mutants (Supp. Fig. 9c,d). Additionally, vertebral bodies (Supp. Fig. 10a,b) were shorter with concomitant severe disruption of growth plates in *Acan*-Cre^+/ERT2^;*gp130*^flfl^ mice compared to *gp130*^fl/fl^ controls (Fig. 7d,e). Analysis of the articular cartilage in *gp130* deleted animals also demonstrated a similar phenotype as found in *Acan*-Cre^+/ERT2^;*Stat3*^fl/fl^ mice, with progressive loss of proteoglycans in both female and male mice, with earlier loss in females (Supp. Fig. 11). These data, which mostly phenocopy the results of *Stat3* deletion in chondrocytes, suggest a gp130/Stat3 signaling circuit that is required for chondrocyte proliferation, prevention of hypertrophy and maintenance of matrix synthesis.

**Figure 7:**
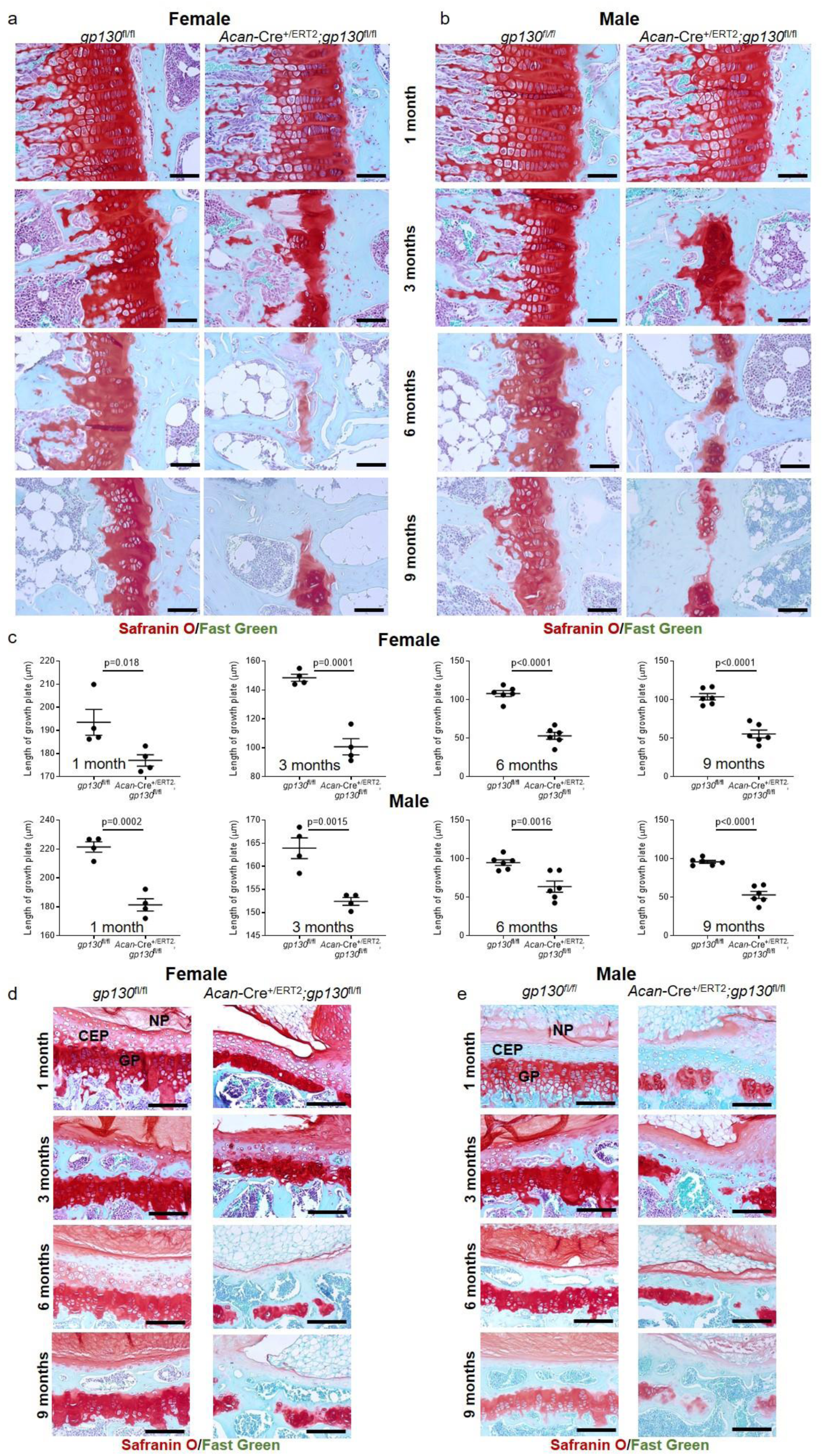
Postnatal loss of *gp130* in chondrocytes phenocopies *Stat3* deletion. Proximal tibial growth plates were significantly reduced in thickness at all ages examined following deletion of *gp130* at P2/P3 in female (a,c) and male (b,c) mice; severe growth plate disruptions were observed at 3 months. Vertebral body heights were reduced, coupled with loss of vertebral growth plates, in both female (d) and male (e) *Acan*-Cre^+/ERT2^;*gp130*^fl/fl^ mice as compared to *gp130*^fl/fl^ controls. AF = annulus fibrosis, CEP = cartilaginous end plate and NP = nucleus pulposus. In all panels, scale bars = 50 µm; n=4-6.

In parallel, we also generated *Acan*-Cre^+/ERT2^;*Lifr*^flfl^ mice, as data from us and others has clearly indicated a role for Lifr/gp130 signaling in skeletal development in mice^11, 58^ and humans^22, 59^. Analysis of Lifr expression in growth plates confirmed a similar expression pattern in the resting and proliferative zones as defined for Stat3 and gp130 (Supp. Fig. 9a). Analysis of growth plates of *Acan*-Cre^+/ERT2^;*Lifr*^flfl^ mice vs. *Lifr^fl/fl^* controls at all ages demonstrated reduced thickness in female and male mice (Fig. 8a-c); at 1 month, the imbalance of the proliferative and hypertrophic zones of the growth plate observed in *Stat3* and *gp130* mutant animals was also present in *Lifr* deleted mice (Supp. Fig. 9). However, in general the skeletal phenotype was milder than *Stat3* and *gp130* deleted mice, as bony bridges of the growth plate were infrequently observed even at 9 months and the articular cartilage, vertebral bodies and growth plates of *Acan*-Cre^+/ERT2^;*Lifr*^fl/fl^ mice were not as severely impacted (Fig. 8d,e; Supp. Figs. 10,11). These results suggest a role for Lifr to mediate IL-6 family cytokine-mediated activation of Stat3 in chondrocytes to promote maintenance of the growth plate, but demonstrate that other receptors are also required.

**Figure 8:**
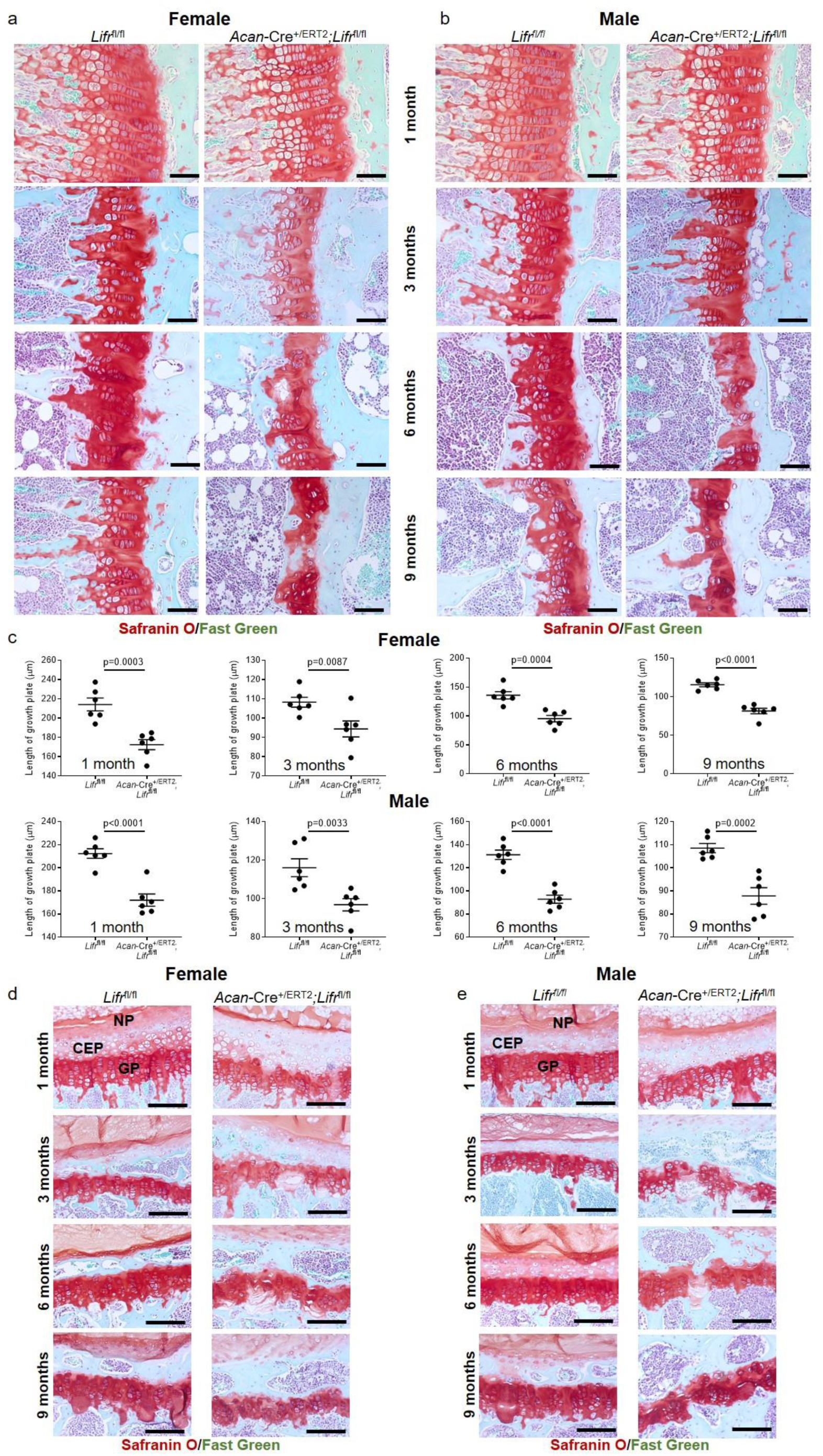
Postnatal loss of *Lifr* in chondrocytes partially phenocopies *Stat3* deletion. Proximal tibial growth plates were significantly reduced in thickness at all ages examined following deletion of *Lifr* at P2/P3 in female (a,c) and male (b,c) mice; growth plate disruptions were not as severe as those observed in *Stat3* and *gp130* deletion mutants. Vertebral growth plates were mildly reduced in thickness, coupled with occasional disruption of integrity at later ages examined, in both female (d) and male (e) *Acan*-Cre^+/ERT2^;*Lifr*^fl/fl^ mice as compared to *Lifr*^fl/fl^ controls. AF = annulus fibrosis, CEP = cartilaginous end plate and NP = nucleus pulposus. In all panels, scale bars = 50 µm; n=6.

### Stat3 overexpression promotes excessive chondrocyte proliferation and rescues growth plate deficiencies observed in *gp130* deleted mice

Postnatal loss of *Stat3* in chondrocytes resulted in premature closure of growth plates accompanied by reduced proliferation and survival of chondrocytes, suggesting this transcription factor is necessary to maintain appropriate mitotic rates. To assess the capacity of Stat3 to elicit a proliferative response, we crossed *Acan*-Cre^+/ERT2^ animals with mice carrying a conditional, constitutively active allele of Stat3 (*Stat3C*) under the control of the *Rosa26* promoter^18^. In these animals, activation of Cre results in excision of a STOP cassette upstream of the *Stat3C* sequence. Analysis of *Acan*-Cre^+/ERT2^;*Rosa26-Stat3C*^fl/+^ mice treated with tamoxifen at P2/P3 and analyzed at 1 month of age revealed dramatic hypercellularity in proximal tibial growth plates (Fig. 9a); hypercellularity was also apparent in the superficial zone and deeper layers in articular cartilage (Fig. 9b). Hyperproliferation in the growth plate was confirmed by EdU incorporation in *Acan*-Cre^+/ERT2^;*Rosa26-Stat3C*^fl/fl^ mice (Fig. 9c), with significantly more EdU^+^ chondrocytes present in Stat3C overexpressing mice as compared to *Rosa26-Stat3C*^fl/fl^ controls. There are also more dividing cells in articular cartilage of Stat3C overexpressing animals at 3 months, although this was not as drammatic as in the growth plate (Supp. Fig. 12). As Stat3 is a known mediator of signaling downstream of gp130 and our data demonstrated similar phenotypes in the growth plates of *Stat3* and *gp130* postnatal deletion mutants, we hypothesized that constitutive activation of Stat3 could rescue the defects in proliferation found in *Acan*-Cre^+/ERT2^;*gp130*^fl/fl^ mice. We generated *Acan*-Cre^+/ERT2^;*gp130*^fl/fl^*Rosa26-Stat3C*^fl/+^ animals and compared their growth plates to *Acan*-Cre^+/ERT2^;*gp130*^fl/fl^ mice at 1 month. In both female and male mice, overexpression of Stat3C in chondrocytes could completely rescue the diminished growth plates in *gp130* deleted mice (Fig. 9d). These data indicate that gp130-mediated activation of Stat3 is required for chondrocyte proliferation, and show that Stat3 is sufficient to promote proliferation in a context-dependent fashion as increased mitosis was most prominent in chondrocytes that normally have the greatest capacity to divide.

**Figure 9:**
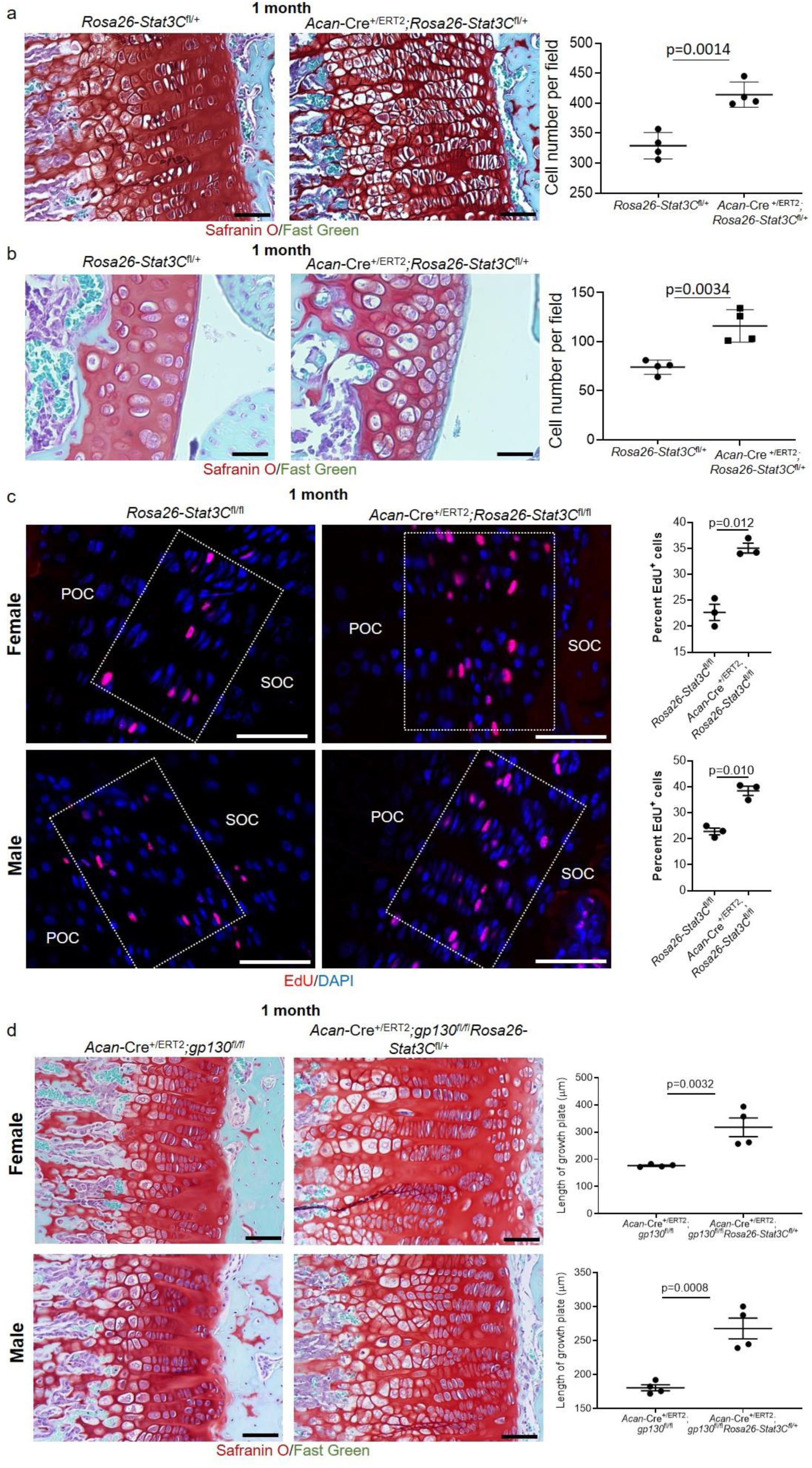
Overexpression of constitutively active Stat3 (Stat3C) in chondrocytes promotes precocious proliferation *in vivo* and rescues growth plate defects in *gp130* deletion mutants. Induction of Stat3C expression at P2/P3 resulted in significant increases in chondrocyte number in both growth plate (a) and articular cartilage (b) in *Acan*-Cre^+/ERT2^;*Rosa26*-Stat3C^fl/+^ mice as compared to *Rosa26*-Stat3C^fl/+^ control animals at 1 month. (c) EdU was injected intraperitoneally 4 hours before harvesting knee joints at 1 month of age. EdU^+^ cells in proximal tibial growth plate chondrocytes were scored. POC = primary ossification center; SOC = secondary ossification center. Boxed areas delineate the growth plate. (d) Induction of constitutively active Stat3 in *Acan*-Cre^+/ERT2^;*gp130*^fl/fl^ mice resulted in significantly thicker growth plates at 1 month of age. In all panels, scale bars = 50 µm; n=4-6 for each group.

### Estrogen increases Stat3 expression and activation in chondrocytes

A relationship between sex hormones and chondrogenesis is well established. In humans, estrogen is responsible in both females and males for accelerating growth at puberty and eventually driving the closure of growth plates; females tend to begin the pubertal growth spurt earlier than males and have higher levels of estradiol at this age (reviewed in^60^). Based on the dimorphic difference in body size at 3 months in *Acan*-Cre^ERT2^;*Stat3*^fl/fl^ females and males, we hypothesized that estrogen may regulate Stat3 levels and/or activity. To address this, we isolated porcine articular chondrocytes from female and male animals and treated them with different levels of estradiol (E2) *in vitro*. These data demonstrated that both female (Fig. 10a) and male (Fig. 10b) articular chondrocytes upregulated STAT3 proteins levels and activity in response to E2, though female chondrocytes demonstrated a maximal response at 0.1 µM while male chondrocytes responded in a dose-dependent manner. To assess the impact of estrogen levels on Stat3 *in vivo*, the clinically-used small molecule aromatase inhibitor Letrozole (which results in lower estradiol levels) was injected intra-peritoneally daily for 7 days into sexually mature female mice. Analysis of sternal chondrocytes revealed that Letrozole reduced both Stat3 levels and activity. Taken together, these data suggest that the enhanced estrogen levels present in growing female mice lead to a differential requirement for Stat3 to enhance skeletal growth.

**Figure 10:**
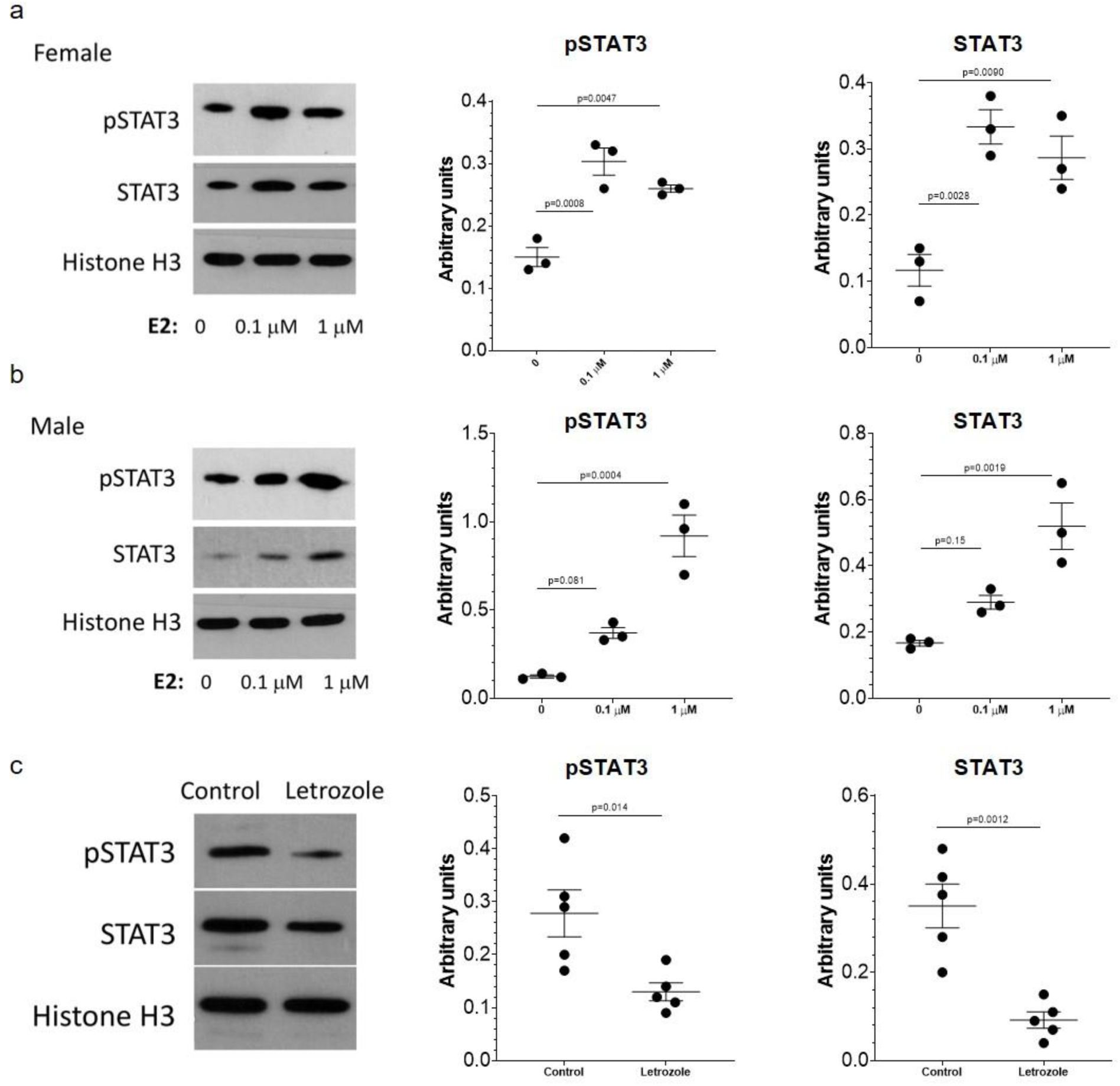
Estrogen influences Stat3 levels and activity *in vitro* and *in vivo*. Addition of estradiol (E2) to female (a) and male (b) pig chondrocytes resulted in dose-dependent increases in Stat3 levels and activity; female chondrocytes are dose-limited in their response to E2 as evidenced by similar induction at the 0.1 and 1.0 µM doses. n = 3. (c) Intra-peritoneal Injection of Letrozole, an aromatase inhibitor that blocks the production of estrogen, reduced Stat3 levels and activity in female mouse sternal chondrocytes after 7 daily injections. n = 5 mice per group.

## Discussion

Here we show that Stat3 is a critical factor controlling the proliferation, survival, maturation and matrix production of chondrocytes in growth plates and articular cartilage. Animals lacking *Stat3* in chondrocytes following postnatal deletion are proportionately smaller than control animals and evidence significantly reduced EdU incorporation, coupled with thinner proliferative zones and increased apoptosis; by 9 months of age, all mice had abnormal growth plate fusions in both appendicular and axial skeletal elements, with increased frequency of fusions found in females. Articular cartilage was also affected, displaying a mild degenerative and pre-hypertrophic phenotype. Females tended to have a more severe phenotype than males, with a more reduced body size than males at 3 months, earlier and more frequent fusions in the growth plate and accelerated changes in the articular cartilage. *Stat3* null chondrocytes were enriched for expression of genes previously associated with maturation in the growth plate and concurrently evidenced diminished expression of genes associated with immature, proliferative chondrocytes. We also showed that deletion of the obligate receptor for IL-6 family cytokines gp130 in chondrocytes postnatally phenocopied loss of *Stat3*, nominating for the first time a role for gp130-Stat3 signaling in regulating immature chondrocyte maintenance; deletion of *Lifr* resulted in a much milder phenotype, potentially due to compensation by other alpha-receptors. Expression of a constitutively active form of Stat3 from the *Rosa26* locus led to excessive proliferation of chondroprogenitors and chondrocytes and rescued the diminutive growth plates found in *gp130* deleted mice, demonstrating Stat3 is both necessary and sufficient to promote cell division in permissive chondrogenic cells. Finally, we showed that estradiol can regulate Stat3 levels and activity *in vitro* and *in vivo*, identifying a potential mechanism of growth acceleration in females during longitudinal bone growth. Together, these data unveil IL-6 family cytokine signaling via gp130-Stat3 as a major regulator of postnatal chondrocyte biology.

Regulation of postnatal bone growth via endochondral ossification has been investigated by several groups using transgenic mice. Deletion of *Sox9*, the master regulator of chondrogenesis, at early postnatal stages resulted in death of the animals^61^; however, deletion at P18 and later using *Acan*-Cre^ERT2^ produced mice with similar phenotypes to those we report here including growth plate shortening and fusions concomitant with reduced proliferation and increased apoptosis, compressed vertebral body heights and pre-arthritic changes in articular cartilage^61^. Some aspects of the phenotype they observed were more severe than in *Stat3* deleted mice, including very rapid loss of matrix secretion proximate to deletion of *Sox9*. Interestingly, Hall et al. demonstrated that STAT3 could bind to the *Sox9* promoter and activate transcription, which may explain some of the phenotypic overlaps^14^. Indeed, deletion of *Stat3* greatly reduced expression of Sox9 in the articular cartilage (Supp. Fig. 8). Moreover, deletion of *Sox9* using *Acan*-Cre^ERT2^ in mice 3 months of age resulted in loss of verterbral endplate integrity coupled with growth plate fusions followed by increases in fibrosis and apoptosis in the nucleus pulposus^42^; these data are strikingly similar to the phenotypes reported here and reinforce the concept of a gene regulatory loop involving *Stat3* and *Sox9* that controls chondroprogenitor output. Although not inducible, another transgenic mouse that evidenced growth plate fusions and mild dwarfism was created by Sims et al., in which the intracellular motifs of gp130 that interact with Stat3 were deleted^62^. By 4 months of age, these mice had femoral lengths an average of 20% less than wild type controls, frequent growth plate fusions and significantly reduced proliferative zones in the growth plate. The authors did not note any vertebral defects, nor whether there were differences between males and females. Furthermore, given the constitutive nature of this knock-in model, it coud not be determined whether the effects on cartilage were direct, and at what developmental stage of chondrogenesis the gp130/Stat3 circuit might be required. The data presented here provide clear evidence that gp130/Stat3 signaling is required postnatally in chondrocytes to coordinate proliferation and differentiation.

Our data from *Stat3* deleted mice reveal a sexually dimorphic response. Stat3 has been implicated in sex-specific responses to ischemia in both the heart^63^ and brain^64^, with increases in Stat3 phosphorylation directly mediated by stimulation with estradiol. The data presented here suggest that estradiol may increase Stat3 levels and activity during the pubertal growth spurt; estradiol levels spike in female C57/B6 mice at 28-30 days^65^. In uninjured brains, Di Domenico et al. identified several differences in female vs. male neurons that depended on Stat3 using a proteomics approach; many of these proteins have been associated with cellular metabolism^66^, consistent with known functions of Stat3 in mitochondrial^67^ and glycolytic metabolism^68^. Stat5a and Stat5b, other members of the same transcription factor family, have demonstrated roles in regulating sexually dimorphic gene expression downstream of growth hormone in the liver^69^. Together, these data support the feasibility of a sex-specific role for Stat3 in enhancing skeletal growth and potentially maintenance that should be explored further.

Growth of the long bones and spine depend upon regulated output from chondroprogenitor cells located in growth plates. In humans, these structures support growth throughout childhood and adolescence, at which point increases in sex hormones are partially responsible for terminal differentiation of chondroprogenitors and closure of growth plates^70^. Abnormalities in the output from chondroprogenitors often manifest in visible alterations in skeletal size. Gigantism can occur when too much growth hormone is present during childhood and adolescence, resulting in excessive linear growth; this condition is treated clinically with a peptide that mimics somatotropin, a hormone that represses growth hormone secretion^71^. In contrast, chondrodysplasias that result in below average stature can result from a myriad of inherited or developed conditions. Mutations in the *FGFR3* gene that produce constitutively active protein are associated with several skeletal conditions including achondroplasia and hypochondroplasia^72^, while inactivating mutations in *LIFR* or *IL6ST* (encodes gp130) genes cause Stuve-Wiedemann syndrome^59, 73^. Moreover, short stature is frequently found in children with chronic inflammatory conditions such as Crohn’s disease and ulcerative colitis which are mediated by excessive levels of pro-inflammatory cytokines including IL-6^74^. Here we showed that *Lifr* deletion specifically in chondrocytes evidenced a noticeably milder phenotype than *Acan*-Cre^ERT2^-mediated deletion of *Stat3* or *gp130*, as well as a much weaker skeletal phenotype than seen in global, constitutive deletion of *Lifr* in mice^58^ or germline mutation in humans. We hypothesize that oncostatin M receptor (Osmr), through which Lif can also signal after heterodimerization with gp130, can compensate for *Lifr* deletion in chondrocytes. Our data from human ontogeny show abundant expression of *OSMR* at relevant stages of human ontogeny (Supp. Fig. 1). Also, Lifr has clear functions in osteoblasts and osteoclasts^58^, and these functions outside the chondrogenic lineage could impact skeletal development. Finally, Stat3 is downstream of growth hormone receptor (GHR), FGFR3 and epidermal growth factor receptor (EGFR) activation as well as IL-6/gp130 signaling, reinforcing the relevance of the current work showing that the correct dosage of Stat3 activity is essential for regulating immature chondrocyte cell fate.

Classically, IL-6/gp130/STAT3 signaling has been viewed as a negative regulator of chondrocyte biology and function and is broadly discussed in the context of arthritis and other inflammatory diseases. IL-6 signaling through IL-6R/gp130 suppresses chondrocyte proliferation^75^, promotes mineralization in articular cartilage^76^, downregulates expression of matrix proteins and increases expression of matrix-degrading proteases^77^. Moreover, blockade of IL-6 *in vivo* in mouse models of osteoarthritis (OA) has been shown to be chondroprotective^78, 79^. Importantly, higher serum levels of IL-6 have been correlated with the development of OA in humans, highlighting the potential for systemic inflammation levels to contribute to local disease; a monoclonal antibody against IL-6R is currently in Phase III clinical trials for treatment of hand OA (NCT02477059). Furthermore, inhibition of Stat3 downstream of exogenous IL-6 is chondroprotective, reducing the severity of OA-like pathology in a mouse model^78^. However, analysis of phenotypes associated with *Stat3*, *gp130* and *Lifr* deletion in chondrocytes during homeostasis reveal essential roles for this pathway in chondroprogenitor proliferation and maintenance. This is reinforced by analysis of patients diagnosed with juvenile arthritis treated with the monoclonal anti-IL-6R antibody Tocilizumab; many of these individuals achieved normalized growth rates when their serum IL-6 levels normalized^80^. It remains to be determined if the Stat3 targets and overall functions in decease and development are identical, or if these effects are context specific with the final outputs determined by signaling interactions with other pathways which are clearly dissimilar in normal growth and diseases. Our results demonstrating the essential role of calibrated gp130/Stat3 signaling in developing chondrocytes suggest that modulation of this axis could have therapeutic benefit in other conditions where growth plate and/or articular cartilage maintenance is impacted.

## Acknowledgments

The authors thank the Molecular Imaging Center core in the Department of Radiology, Keck School of Medicine, University of Southern California and Mr. Tautis Skorka for performing high-resolution micro-CT scanning of the samples for this study. We thank Volume Graphics, GmbH (Heidelberg, Germany) for providing complimentary access to VGSTUDIO MAX 3.4 used in this work (https://www.volumegraphics.com). All schematics were created with Biorender.com and we thank the USC Stem Cell Flow Cytometry Core for their assistance and support. This work was supported by National Institutes of Health grant R01AR071734 (DE and KL), the National Institutes of Health grant R01AG058624 (DE), Department of Defence grant W81XWH-13-1-0465 (DE) and the California Institute of Regenerative Medicine grant TRAN1-09288 (DE).

## Data availability

All scRNA-seq data is deposited in GEO and is available under access number GSEXXXX.

## Author contributions

NQL, YL, LL, JL, DG, JZ, TJ, ZB, JM, JT, RS, AS, NL, BVH, KL and DE performed experiments, analyzed data and interpreted results. LW, FAP, KL and DE provided funding support. NQL, BVH, KL and DE designed experiments and conceived the conceptual framework for the study, as well as wrote the manuscript. All authors reviewed the manuscript.

---

**Supplementary Figure 1:**
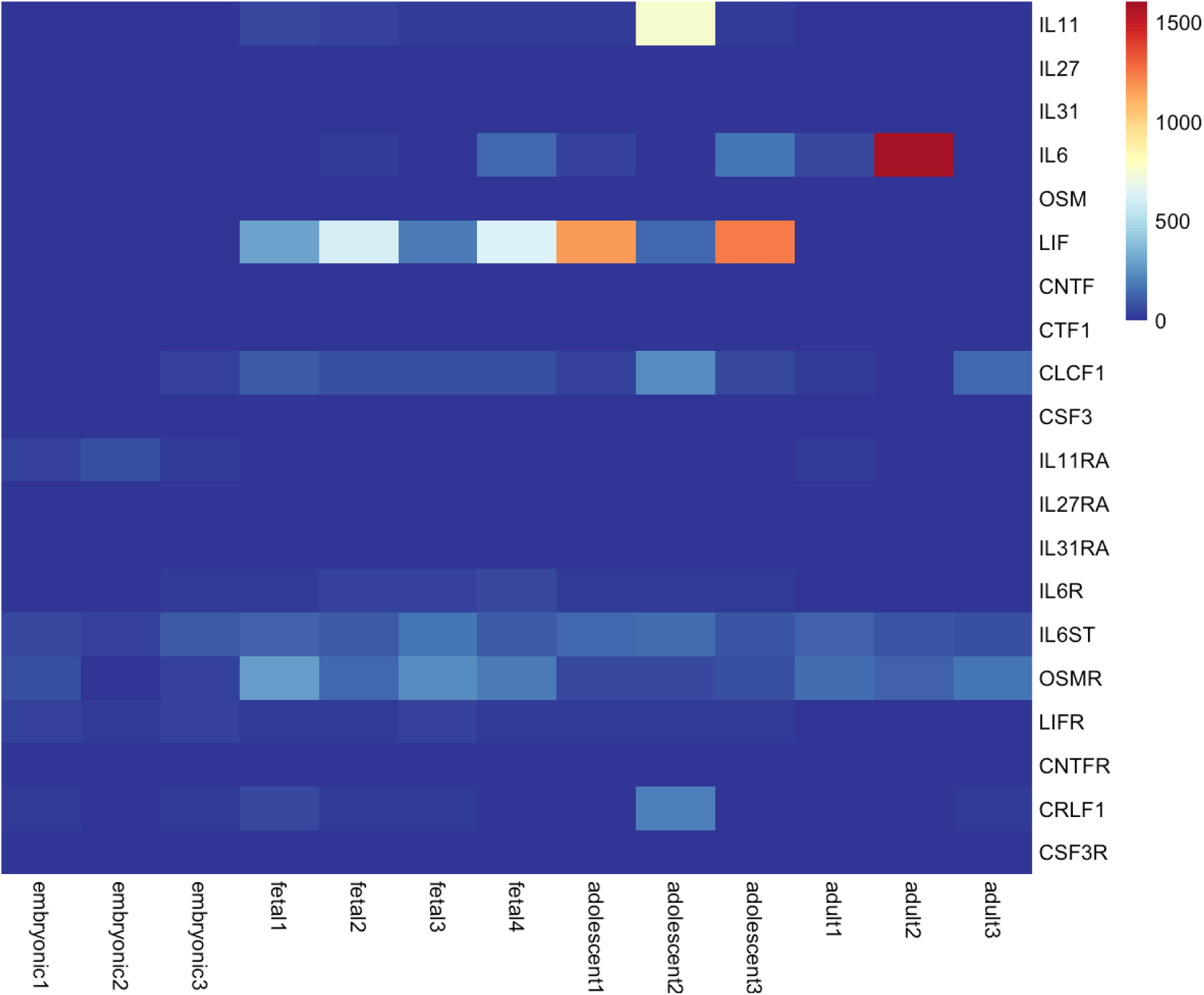
Expression patten of IL-6 family cytokines and receptors during human ontogeny. Analysis of RNA-sequencing data from 4 stages of human ontogeny^12^ demonstrated that gp130 (IL6ST) is expressed at all stages of development, while expression of LIF is enriched at fetal and adolescent stages when growth plates are active in humans. Normalized expression values are shown.

**Supplementary Figure 2:**
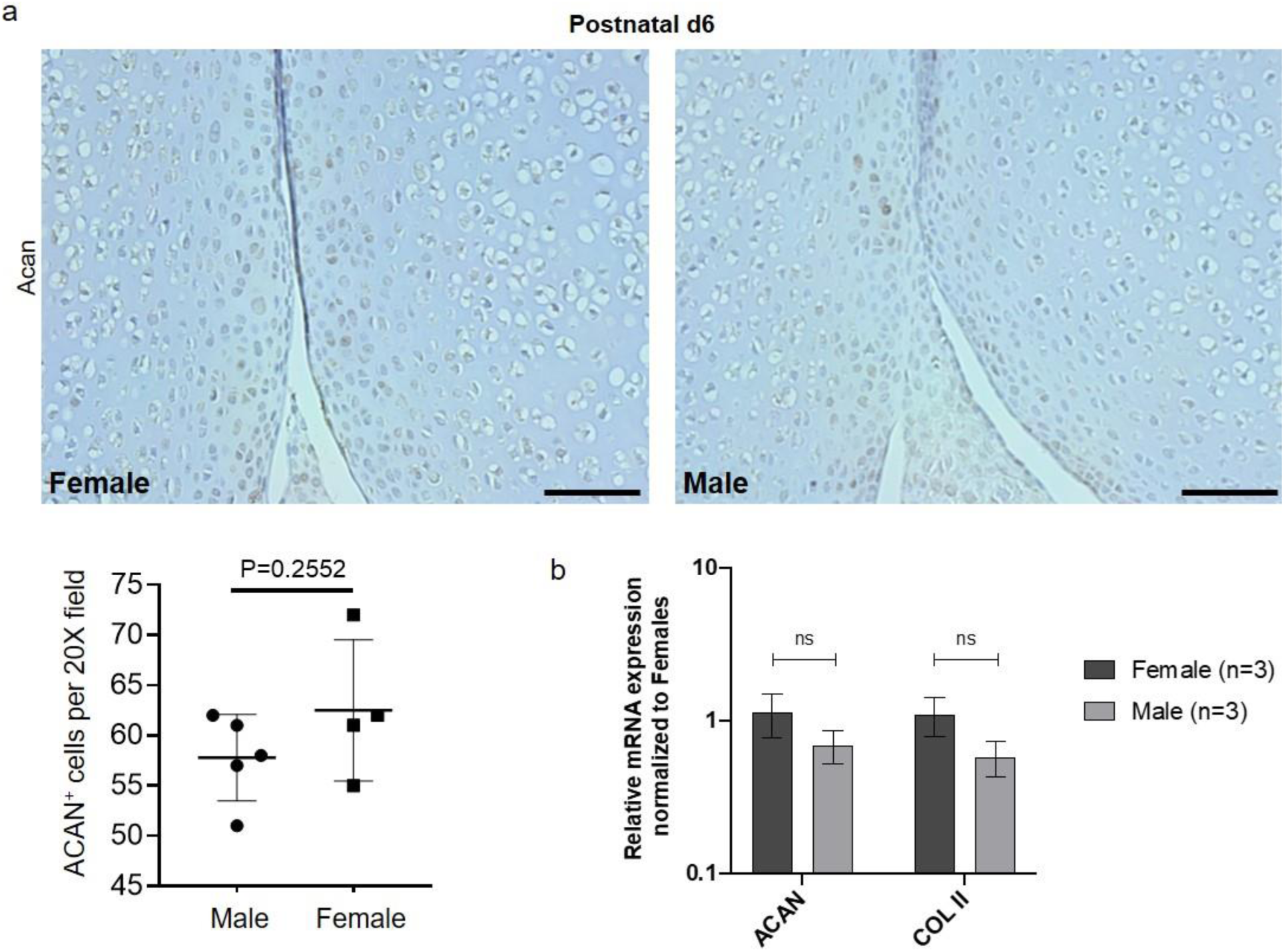
Levels of aggrecan protein and mRNA are not different between females and males at the time of Cre induction. (a) Immunohistochemistry of femoral heads for Acan at postnatal day 6 (P6) demonstrated no significant differences between wild type females and males. n = 5. (b) qPCR for *Acan* and *Col2a1* on chondrocytes isolated from wild type mouse femoral heads showed similar expression levels between females and males. n = 3; scale bars = 50 µm.

**Supplementary Figure 3:**
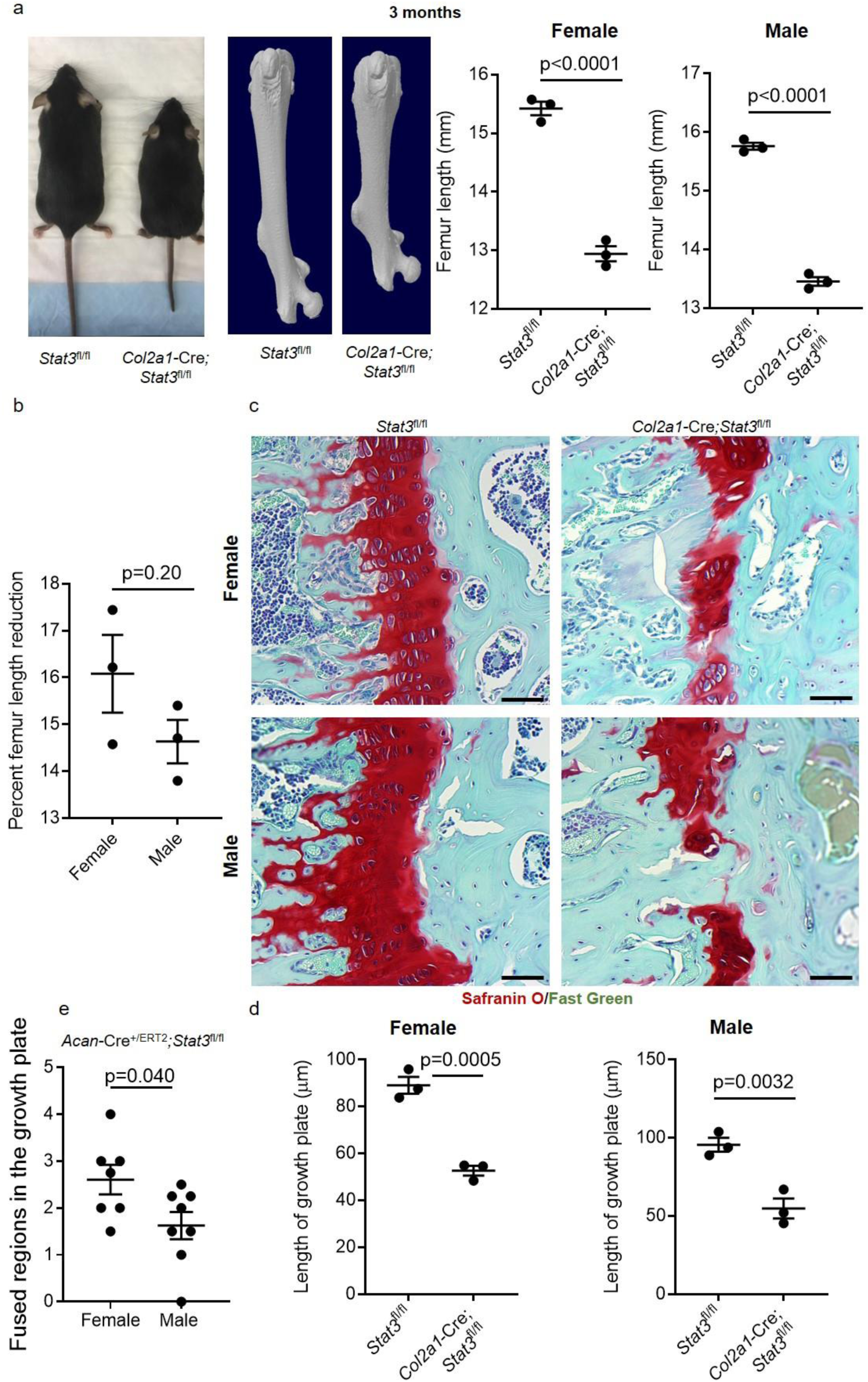
Constitutive deletion of *Stat3* in chondrocytes via *Col2a1*-Cre resulted in reduced body size and growth plate fusions at 3 months. (a) Analysis of control *Stat3*^fl/fl^ and constitutive *Col2a1*-Cre;*Stat3*^fl/fl^ female and male mice at 3 months demonstrated substantially smaller body size and statistically significant reductions in femoral length (b) as determined by microCT. (c) Histological assessment (Safranin O/Fast green staining) of growth plates revealed reduced thickness (d) and fusions in both female and male mice. For all experiments, n = 3; scale bars = 50 µm. (e) *Acan*-Cre^+/ERT2^;*Stat3*^fl/fl^ females evidenced significantly more frequent fusions of the growth plate at 3 months versus *Acan*-Cre^+/ERT2^;*Stat3*^fl/fl^ males; n = 7-8.

**Supplementary Figure 4:**
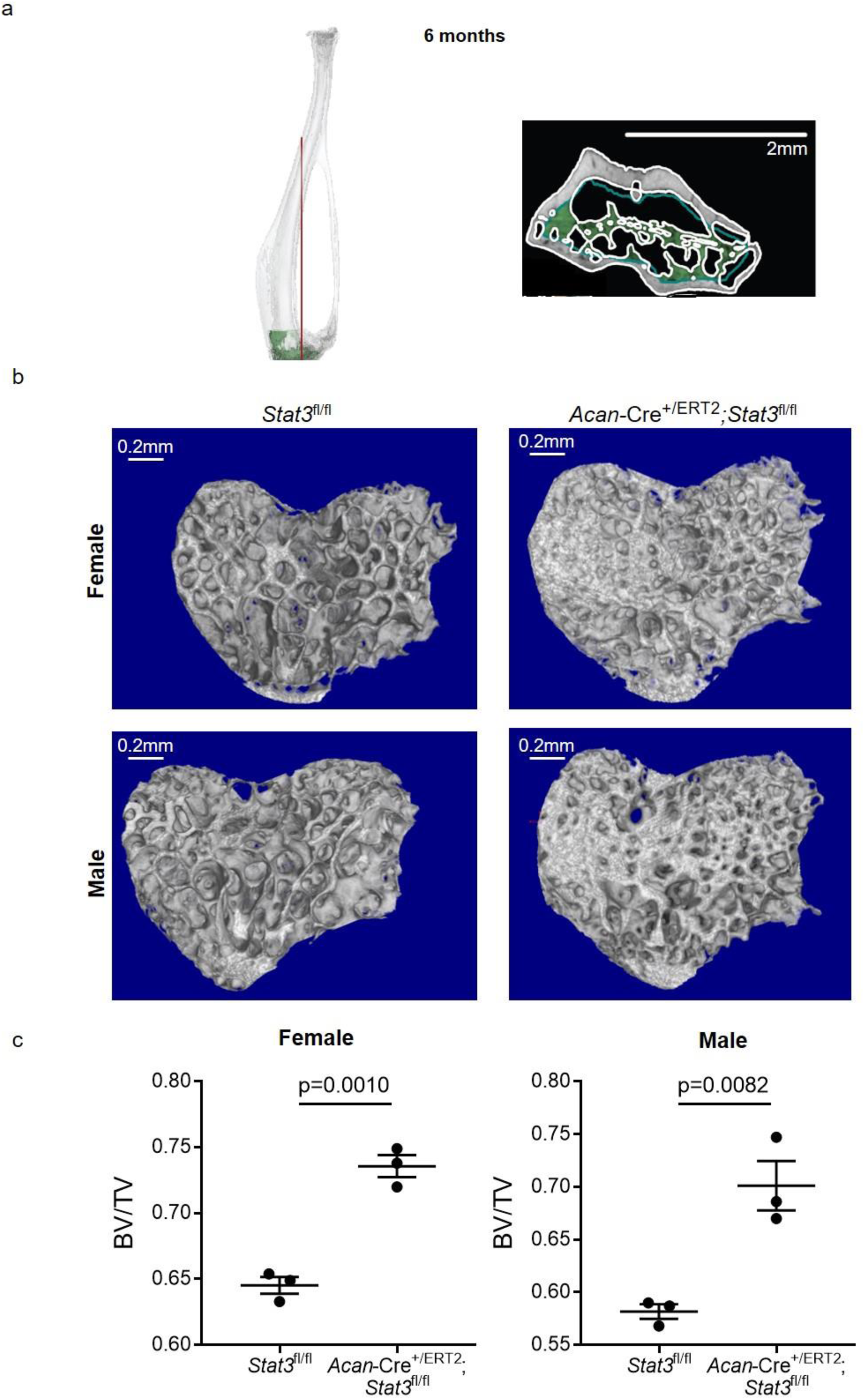
Early postnatal deletion of *Stat3* in chondrocytes increased bone density at 6 months. (a) microCT analysis of control *Stat3*^fl/fl^ and *Acan*-Cre^+/ERT2^;*Stat3*^fl/fl^ mice induced with tamoxifen at P2/P3 revealed loss of *Stat3* increased bone density in the proximal tibial epiphyseal region. The red line on the 3D tibia surface rendering demonstrates the location of transversal cut to visualize bounders between cortical and trabecular bone for region of interest (ROI) segmentation. The 2D green shaded area represents a section of the 3D ROI used to calculate bone volume to total volume (BV/TV). (b) Representative 3D renderings of the ROI in the proximal tibial epiphysis used to calculate BV/TV (c) in for both female and male mice of the indicated genotypes. For all experiments, n = 3.

**Supplementary Figure 5:**
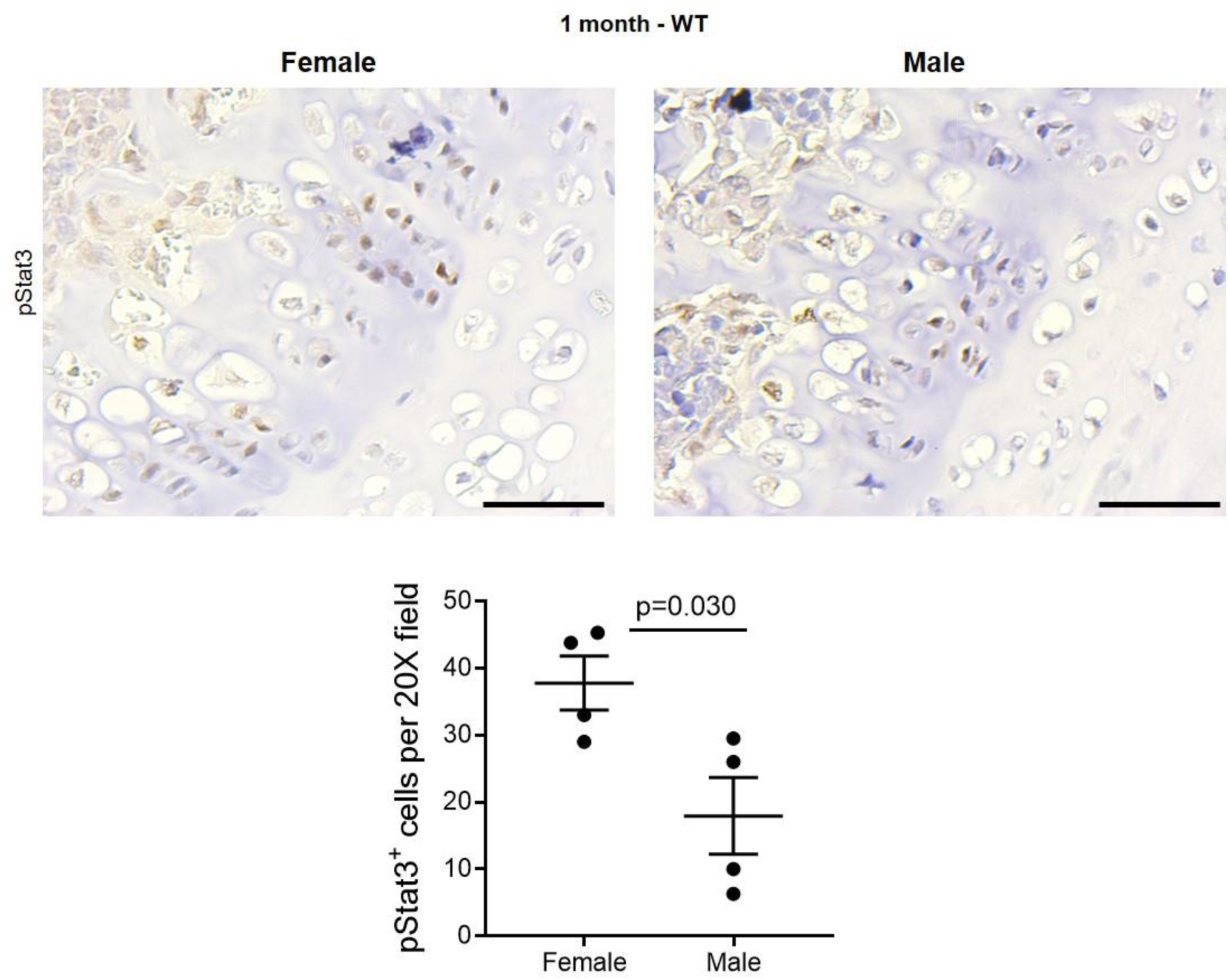
Stat3 activity is enriched in the vertebral growth plates of female versus male wild type mice. Immunohistochemistry for activated Stat3 (pStat3) in vertebral growth plates of wild type female and male mice at 1 month of age demonstrated significantly more active Stat3 in female mice. Note the localization of pStat3 in the proliferative zone. n = 4; scale bars = 50 µm.

**Supplementary Table 2:** Average values of lumbar (L) vertebral body height and endplate areas as determined by microCT. Average value ± standard deviation is shown; n=4 males and females of each genotype pooled. Data were generated based on the measurements described in Supplementary Table 3. VVBH = Ventral (anterior) vertebral body height, DVBH = Dorsal (posterior) vertebral body height, CSA = Cranial endplate surface area, CaSA = Caudal endplate surface area. **Bold** = p <0.001, *italics* = p<0.01, <u>underline</u> = p<0.05 and normal text = not significant.

|  | VVBH (mm) |  | DVBH (mm) |  | CSA (mm<sup>2</sup>) |  | CaSA (mm<sup>2</sup>) |  |
| --- | --- | --- | --- | --- | --- | --- | --- | --- |
|  | Control | *Stat3* deleted | Control | *Stat3* deleted | Control | *Stat3* deleted | Control | *Stat3* deleted |
| L1 | 2.45±0.17 | 1.99±0.07 | 2.89±0.18 | 2.18±0.11 | 1.57±0.06 | 1.90±0.26 | 1.82±0.02 | 2.26±0.42 |
| L2 | 2.77±0.09 | 2.13±0.09 | 3.15±0.10 | 2.37±0.13 | 1.74±0.12 | 1.89±0.19 | <u>2.00±0.09</u> | <u>2.26±0.18</u> |
| L3 | 2.88±0.10 | 2.10±0.03 | 3.37±0.09 | 2.37±0.06 | 1.78±0.10 | 2.10±0.13 | 1.82±0.06 | 2.37±0.26 |
| L4 | 2.88±0.04 | 2.14±0.05 | 3.31±0.08 | 2.33±0.07 | <u>1.82±0.11</u> | <u>2.18±0.18</u> | 2.08±0.09 | 2.51±0.10 |
| L5 | 2.87±0.14 | 2.14±0.10 | 3.18±0.25 | 2.37±0.07 | <u>2.08±0.07</u> | <u>2.40±0.17</u> | <u>2.21±0.12</u> | <u>2.65±0.29</u> |
| L6 | 2.73±0.12 | 2.01±0.10 | 2.96±0.18 | 1.88±0.02 | 2.07±0.07 | 2.38±0.10 | 1.86±0.10 | 2.45±0.16 |

**Supplementary Table 3:** Measurement description on the lumbar vertebrae (L1-L6). Landmark-based data were collected from the lumbosacral region and pelvic girdle. Eight measurements were collected for each lumbar vertebra and then used to generate the data in Supplementary Table 2.

|  | Name | Descriptions | Abbreviation |
| --- | --- | --- | --- |
| 1 | Ventral (anterior) vertebral body height<sup>1</sup> | Vertebral body height, measured from ventral view | VVBH |
| 2 | Dorsal (posterior) vertebral body height<sup>1</sup> | Vertebral body height, measured from dorsal view | DVBH |
| 3 | Cranial transversal diameter of the vertebral body<sup>1</sup> | Transverse (mediolateral) diameter of the vertebral body. Measured on the cranial view from the external cortex of the right border to the external cortex of the left border. | CTD |
| 4 | Cranial DV(AP) diameter of the vertebral body<sup>1</sup> | Dorsoventral (anteroposterior) diameter of the vertebral body. Length measured on the cranial at the midline of the vertebral body from the external cortex of the anterior (ventral) border to the external cortex of the posterior (dorsal) border. | CDVD |
| 5 | Caudal transversal diameter of the vertebral body<sup>1</sup> | Transverse (mediolateral) diameter of the vertebral body. Measured on the caudal view from the external cortex of the right border to the external cortex of the left border. | CaTD |
| 6 | Caudal DV(AP) diameter of the vertebral body<sup>1</sup> | Dorsoventral (anteroposterior) diameter of the vertebral body. Length measured on the caudal at the midline of the vertebral body from the external cortex of the anterior (ventral) border to the external cortex of the posterior (dorsal) border. | CaDVD |
| 7 | Cranial endplate surface area<sup>2</sup> |  | CSA |
| 8 | Caudal endplate surface area<sup>2</sup> |  | CaSA |

<sup>1</sup>measurement in mm; <sup>2</sup>measurement in mm<sup>2</sup>

**Supplementary Figure 6:**
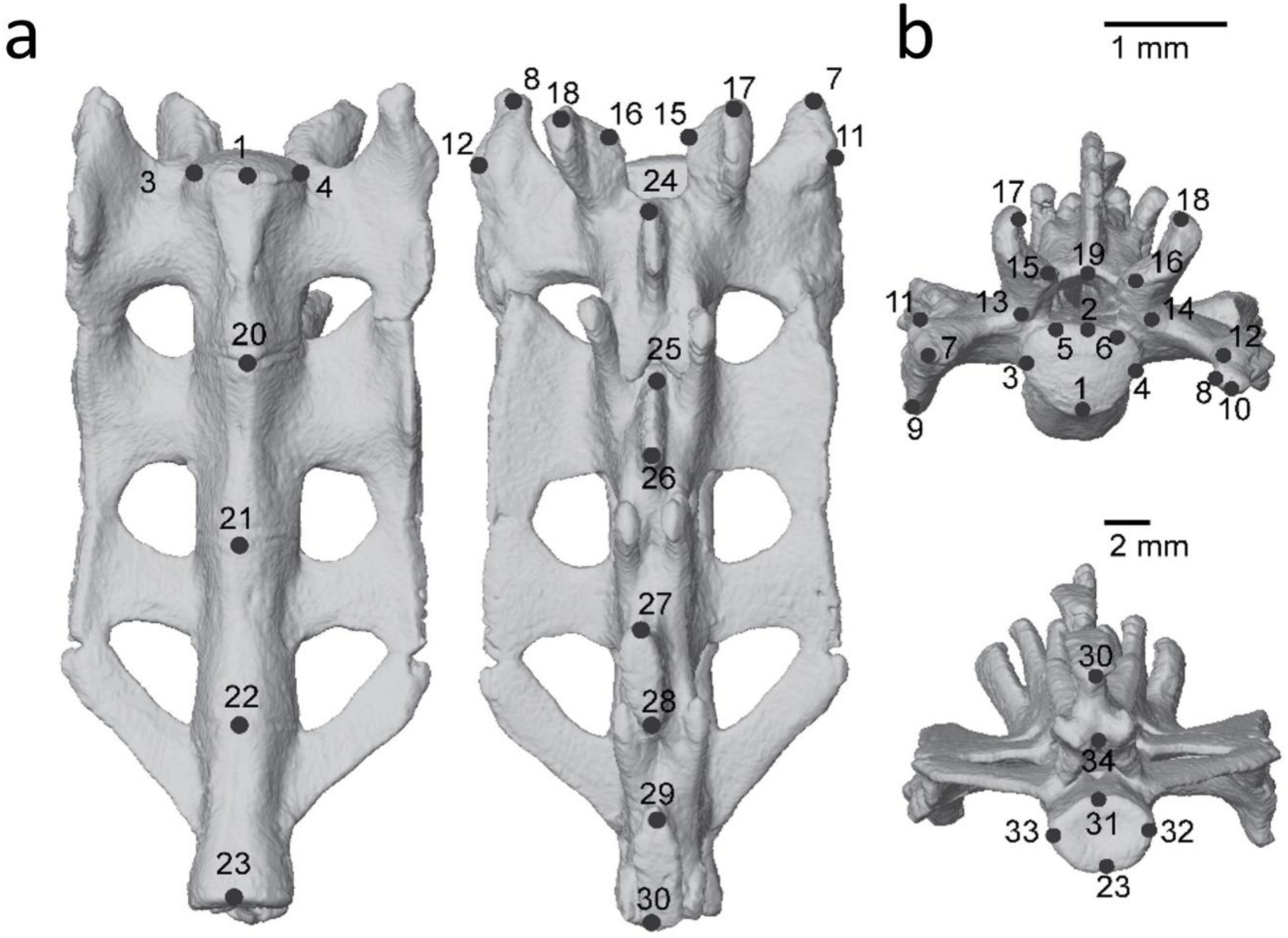
Sacrum of a male *Acan*-Cre^+/ERT2^;*Stat3*^fl/fl^ mouse illustrating the 33 landmarks used as a basis for geometric morphometrics. (a) Ventral and dorsal views. (b) Cranial and caudal views. Landmarks are described in the Materials and methods section. Imaging was conducted at 6 months.

**Supplementary Figure 7:**
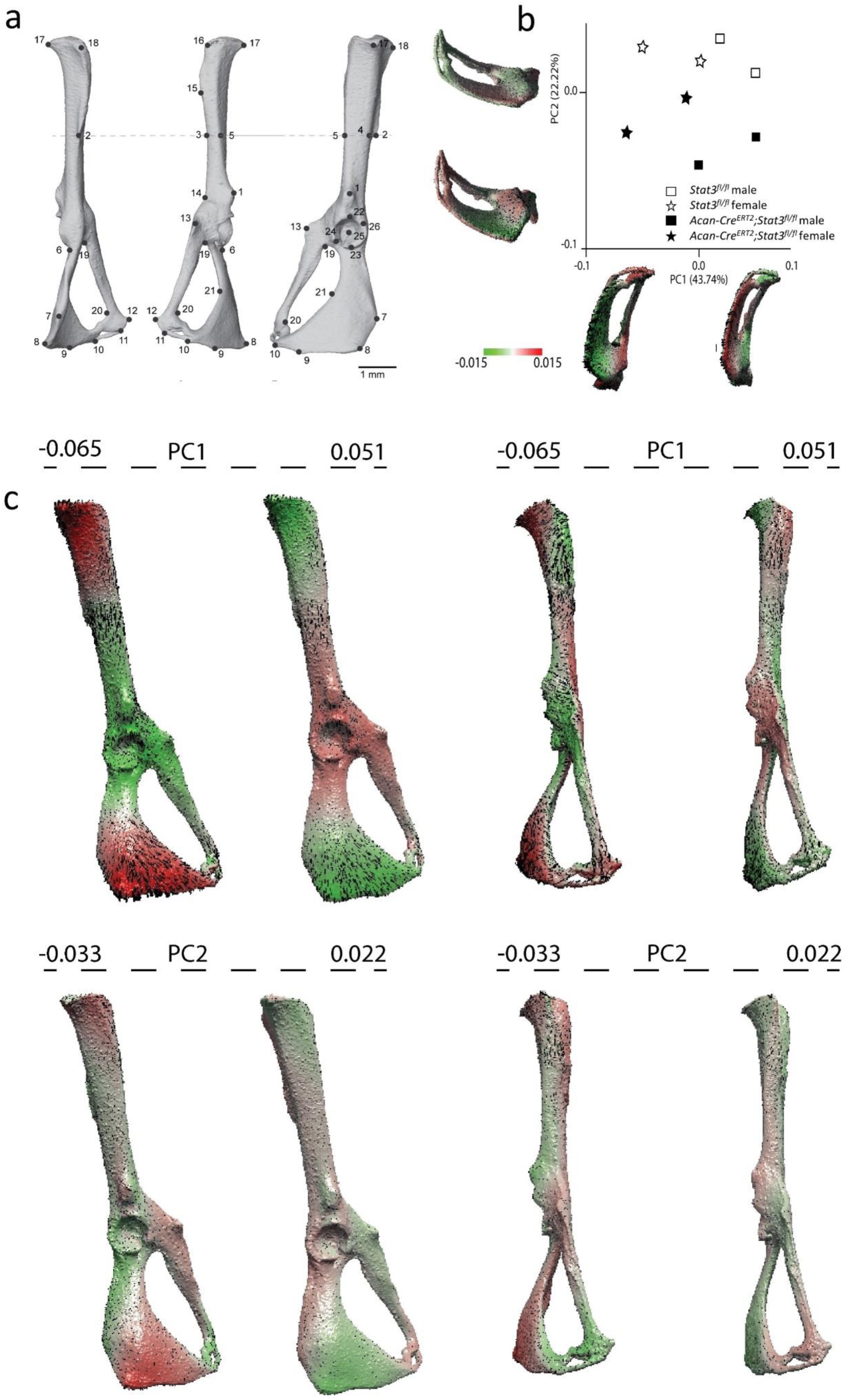
Significant skeletal structural changes result after deletion of *Stat3*. (a) Left pelvis of a male *Acan*-Cre^+/ERT2^;*Stat3*^fl/fl^ mouse in dorsal, ventral and lateral (left to right) views illustrating the 26 landmarks used as a basis for geometric morphometrics. (a) Ventral and dorsal views. (b) Principal component analysis (PCA) of pelvises from Stat3 deleted and control mice based on distances between 26 anatomical landmarks. Animals clearly segregate based on genotype. (c) Deviations along PC1 (above) and PC2 (below) are shown on 3D surface maps of control (left) and Stat3 deleted (right) male pelvic regions. Maps are colored based on degree of deformation with respect to PC values, with red representing features more prominent in control and green more prominent in Stat3 deleted animals, respectively.

**Supplementary Figure 8:**
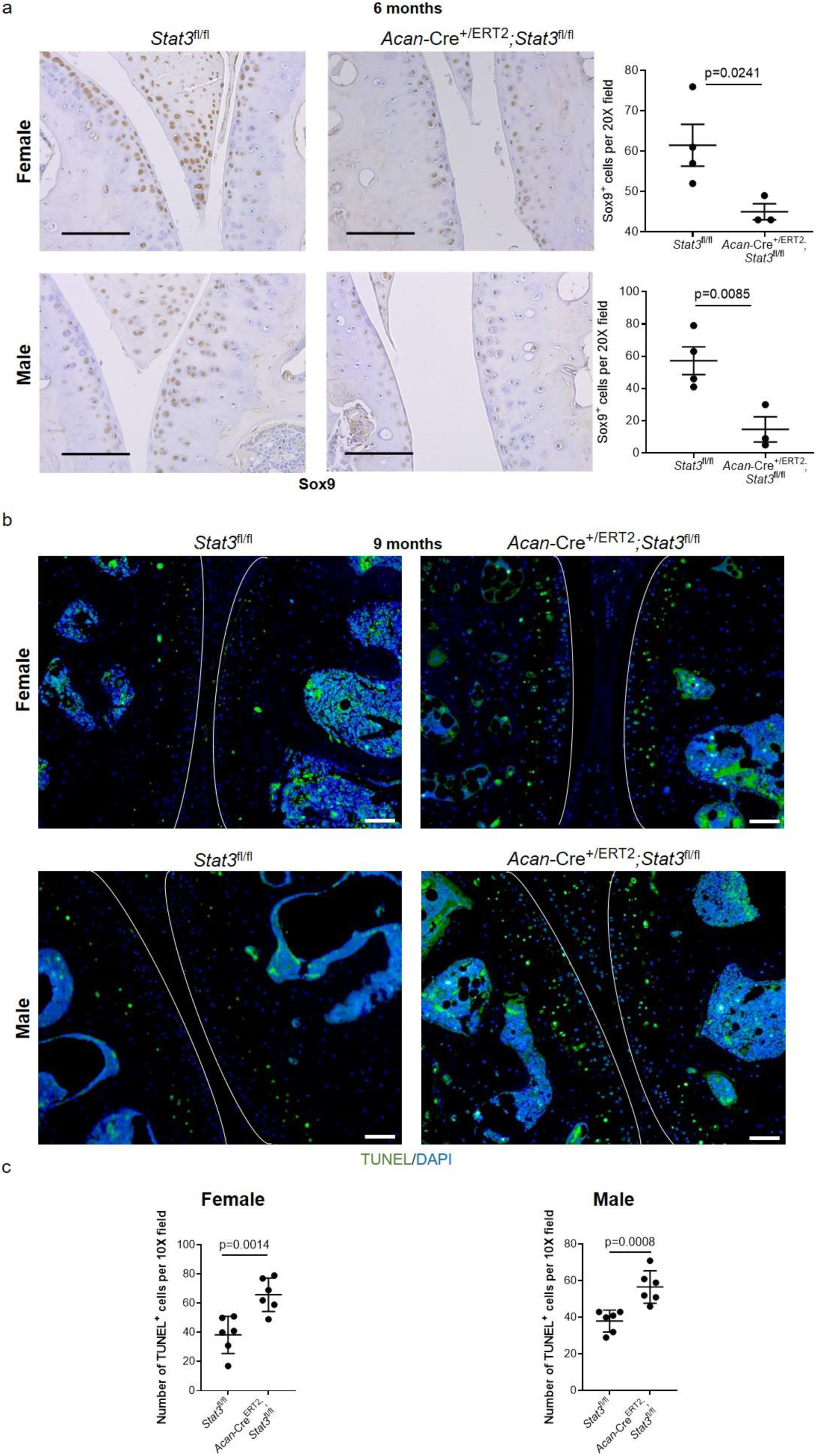
Deletion of *Stat3* results in reduced Sox9 expression and increased apoptosis in articular cartilage. (a) Following tamoxifen administration at P2/3, animals were sacrificed at 6 months. Note reduced frequency and intensity of Sox9 staining in *Acan*-Cre^+/ERT2^;*Stat3*^fl/fl^ animals. Representative images are shown and scale bars = 50 µm; n = 3-4. (b) Assessment of apoptosis in articular cartilage 9 months after *Stat3* deletion by TUNEL staining demonstrated increased cell death (c) in *Acan*-Cre^+/ERT2^;*Stat3*^fl/fl^ animals. The articular surface is delineated by white lines in (b). n = 6; scale bars = 50 µm.

**Supplementary Figure 9:**
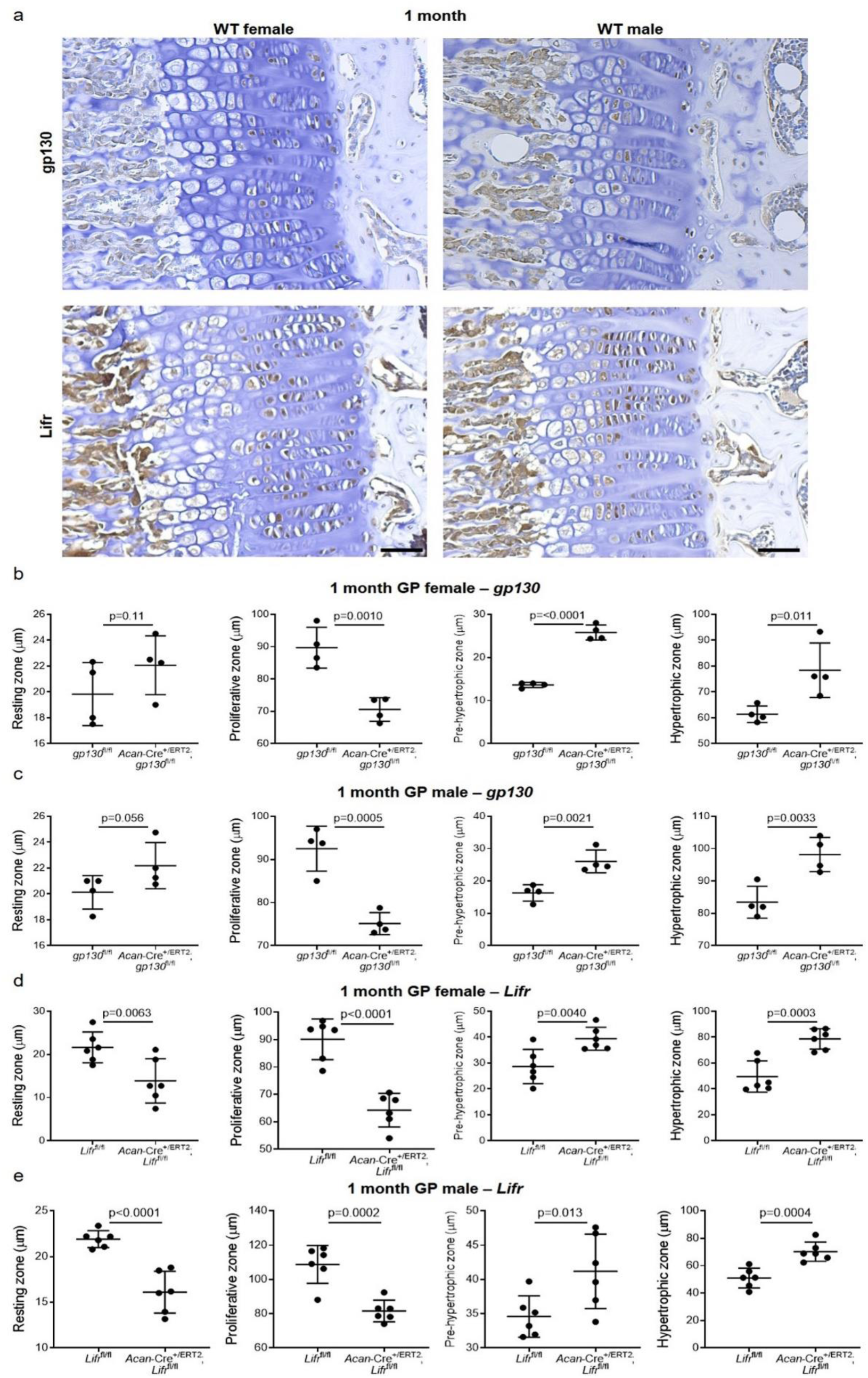
Deletion of *gp130* and *Lifr* in chondrocytes resulted in reduced proliferation and increased hypertrophy in the growth plate at 1 month as well as growth plate fusions. (a) Localization of gp130 and Lifr expression in growth plate chondrocytes in wild type mice. Note the enrichment of both receptors in the proliferative zone. n = 3, representative images are shown; scale bars = 50 µm. (b) *Acan*-Cre^+/ERT2^;*gp130*^fl/fl^ females evidenced significantly more frequent fusions of the growth plate at 3 months versus *Acan*-Cre^+/ERT2^;*gp130*^fl/fl^ males; n = 4. (c-f) Measurement of each zone of the growth plate demonstrated significant reductions in the proliferative zone in both females and males of *Acan*-Cre^+/ERT2^;*gp130*^fl/fl^ and *Acan*-Cre^+/ERT2^;*Lifr*^fl/fl^ animals as compared to controls; in contrast, both the pre-hypertrophic and hypertrophic zones increased in size in both sexes of *gp130* and *<u>Lifr</u>* animals vs. controls. n = 4-6.

**Supplementary Figure 10:**
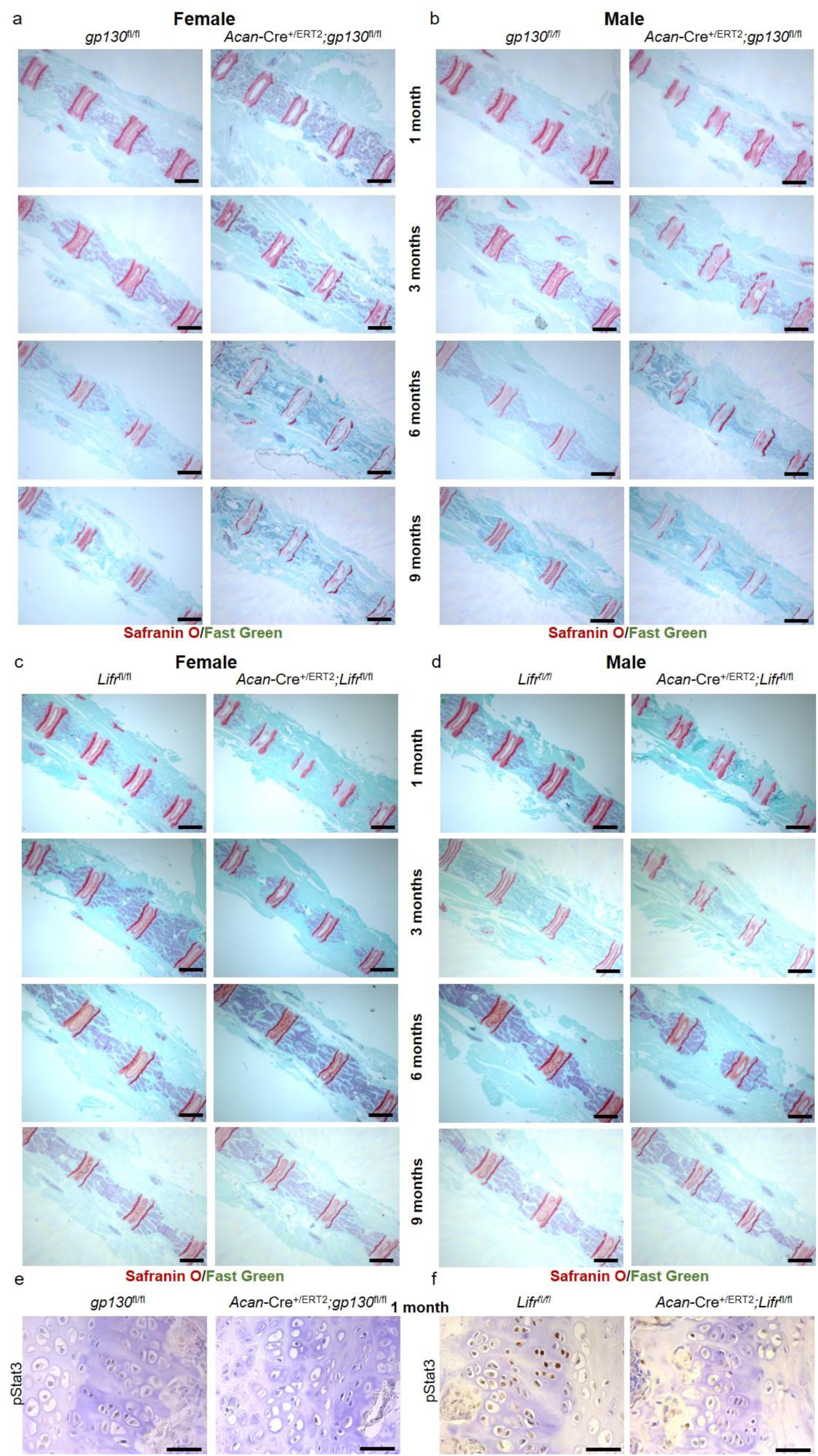
Postnatal deletion of *gp130*, and to a lesser extent *Lifr*, affects vertebral growth plates and bodies. (a) Coronal sections of caudal vertebrae of *Acan*-Cre^ERT2^;*gp130*^fl/fl^ and control *gp130*^fl/fl^ mice demonstrate shorter vertebral bodies in both female and (b) male mice. Note the increased number of vertebrae visible in each image in the *gp130* deleted mice vs. controls, as well as the degeneration of the growth plates. Deletion of *Lifr* resulted in a milder phenotype, with moderate compression of the caudal vertebrae in both females (c) and males (d); degeneration of the growth plates is not obvious until the 9 month timepoint. In a-d, scale bars = 1 mm. (e) Deletion of both *gp130* and *Lifr* (f) resulted in substantial loss of Stat3 activity (pStat3) in vertebral growth plates at 1 month; scale bars = 50 µm. For all panels, n = 4. Representative images are shown.

**Supplementary Figure 11:**
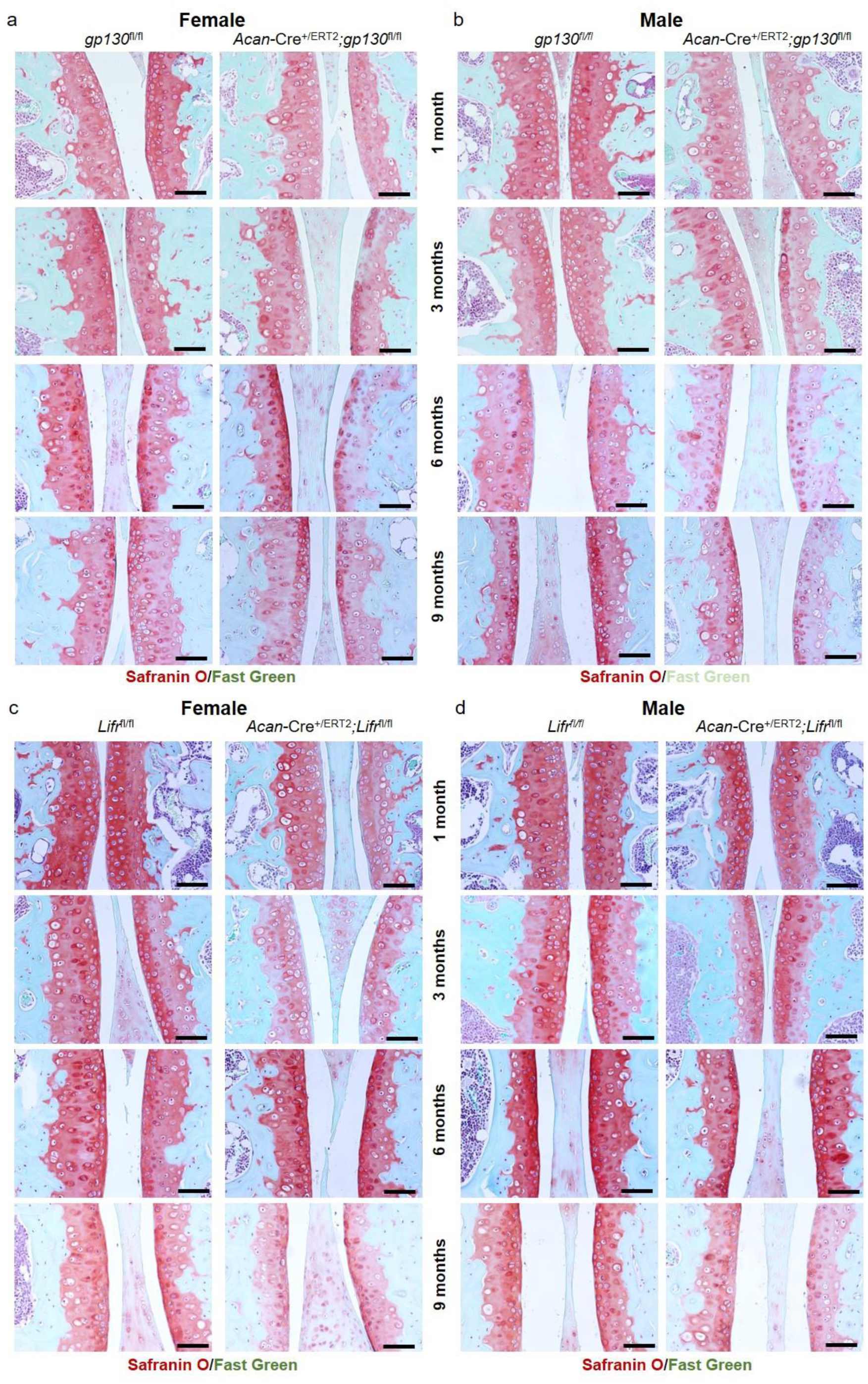
Postnatal *gp130*, and to a lesser extent *Lifr*, deletion results in subtle changes in articular cartilage of the knee joint. (a) Administration of tamoxifen at P2/P3 to *Acan*-Cre^ERT2^;*gp130*^fl/fl^ and control *gp130*^fl/fl^ female and (b) male mice elicited reduced proteoglycan staining in articular cartilage at 6 and 9 months of age in both sexes. Deletion of *Lifr* resulted in a milder and delayed phenotype, with proteoglycan depletion not evident until 9 months of age in both females (c) and males (d). For all panels, representative images are shown; n = 4 and scale bars = 50 µm.

**Supplementary Figure 12:**
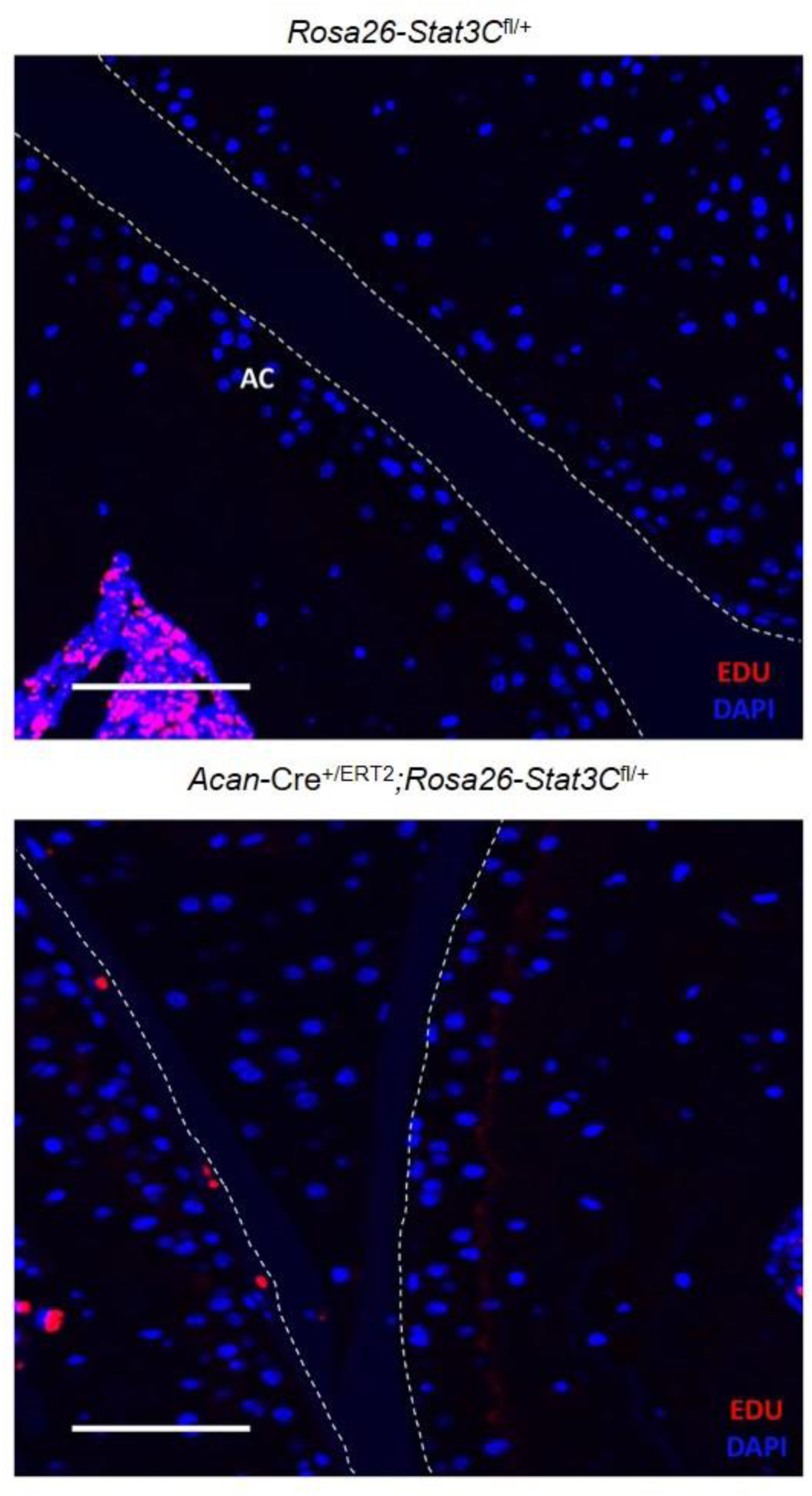
Increased proliferation in articular cartilage following overexpression of constitutively active Stat3. EdU was injected intraperitoneally each day for 4 days before harvesting knee joints at 1 month of age. Representative images of the articular cartilage (AC) are shown. Scale bars = 50 µm; n=5 for each group.

## Notes

### Competing Interest Statement

The authors have declared no competing interest.

## References

1. Nilsson, O. & Baron, J. Fundamental limits on longitudinal bone growth: growth plate senescence and epiphyseal fusion. Trends in endocrinology and metabolism: TEM 15, 370–374 (2004).

2. Mackie, E.J., Ahmed, Y.A., Tatarczuch, L., Chen, K.S. & Mirams, M. Endochondral ossification: How cartilage is converted into bone in the developing skeleton. The International Journal of Biochemistry & Cell Biology 40, 46–62 (2008).

3. Mizoguchi, I., Nakamura, M., Takahashi, I., Kagayama, M. & Mitani, H. An immunohistochemical study of localization of type I and type II collagens in mandibular condylar cartilage compared with tibial growth plate. Histochemistry 93, 593–599 (1990).

4. Anderson, H.C. et al. Impaired Calcification Around Matrix Vesicles of Growth Plate and Bone in Alkaline Phosphatase-Deficient Mice. The American Journal of Pathology 164, 841–847 (2004).

5. Yang, L., Tsang, K.Y., Tang, H.C., Chan, D. & Cheah, K.S. Hypertrophic chondrocytes can become osteoblasts and osteocytes in endochondral bone formation. Proceedings of the National Academy of Sciences 111, 12097–12102 (2014).

6. Yu, Y.Y., Lieu, S., Lu, C. & Colnot, C. Bone morphogenetic protein 2 stimulates endochondral ossification by regulating periosteal cell fate during bone repair. Bone 47, 65–73 (2010).

7. Howell, F.R., Mahood, J.K. & Dickson, R.A. Growth beyond skeletal maturity. Spine 17, 437–440 (1992).

8. Crowder, C. & Austin, D. Age ranges of epiphyseal fusion in the distal tibia and fibula of contemporary males and females. Journal of forensic sciences 50, 1001–1007 (2005).

9. Newton, P.T. et al. A radical switch in clonality reveals a stem cell niche in the epiphyseal growth plate. Nature 567, 234–238 (2019).

10. Kimura, H., Ng, J.M.Y. & Curran, T. Transient Inhibition of the Hedgehog Pathway in Young Mice Causes Permanent Defects in Bone Structure. Cancer Cell 13, 249–260 (2008).

11. Shkhyan, R. et al. Drug-induced modulation of gp130 signalling prevents articular cartilage degeneration and promotes repair. Annals of the Rheumatic Diseases 77, 760 (2018).

12. Ferguson, G.B. et al. Mapping molecular landmarks of human skeletal ontogeny and pluripotent stem cell-derived articular chondrocytes. Nature Communications 9, 3634 (2018).

13. Liu, Y., Sepich, D.S. & Solnica-Krezel, L. Stat3/Cdc25a-dependent cell proliferation promotes embryonic axis extension during zebrafish gastrulation. PLoS genetics 13, e1006564 (2017).

14. Hall, M.D., Murray, C.A., Valdez, M.J. & Perantoni, A.O. Mesoderm-specific Stat3 deletion affects expression of Sox9 yielding Sox9-dependent phenotypes. PLoS genetics 13, e1006610 (2017).

15. Mosley, B. et al. Dual oncostatin M (OSM) receptors. Cloning and characterization of an alternative signaling subunit conferring OSM-specific receptor activation. J Biol Chem 271, 32635–32643 (1996).

16. Henry, S.P. et al. Generation of aggrecan-CreERT2 knockin mice for inducible Cre activity in adult cartilage. Genesis (New York, N.Y. : 2000) 47, 805–814 (2009).

17. Liu, Y., Li, P.-K., Li, C. & Lin, J. Inhibition of STAT3 Signaling Blocks the Anti-apoptotic Activity of IL-6 in Human Liver Cancer Cells. Journal of Biological Chemistry 285, 27429–27439 (2010).

18. Fogli, L.K. et al. T Cell–Derived IL-17 Mediates Epithelial Changes in the Airway and Drives Pulmonary Neutrophilia. The Journal of Immunology 191, 3100 (2013).

19. Farley, F.W., Soriano, P., Steffen, L.S. & Dymecki, S.M. Widespread recombinase expression using FLPeR (flipper) mice. Genesis 28, 106–110 (2000).

20. Ovchinnikov, D.A., Deng, J.M., Ogunrinu, G. & Behringer, R.R. Col2a1-directed expression of Cre recombinase in differentiating chondrocytes in transgenic mice. Genesis 26, 145–146 (2000).

21. Ahn, S. & Joyner, A.L. Dynamic changes in the response of cells to positive hedgehog signaling during mouse limb patterning. Cell 118, 505–516 (2004).

22. Wu, L. et al. Human Developmental Chondrogenesis as a Basis for Engineering Chondrocytes from Pluripotent Stem Cells. Stem Cell Reports 1, 575–589 (2013).

23. Love, M.I., Huber, W. & Anders, S. Moderated estimation of fold change and dispersion for RNA-seq data with DESeq2. Genome Biol 15, 550–550 (2014).

24. Wu, L. et al. Kappa opioid receptor signaling protects cartilage tissue against posttraumatic degeneration. JCI insight 2 (2017).

25. Lebrun, R. in Anthropological Institute & Museum, Vol. Ph.D. Thesis in Paleontology (Université Montpellier II, University of Zürich, Zurich; 2007).

26. Specht, M. in Anthropological Institute & Museum, Vol. Ph.D. (University of Zürich, Zurich; 2007).

27. Specht, M., Lebrun, R. & Zollikofer, C.P.E. Visualizing shape transformation between chimpanzee and human braincases. Visual Comput 23, 743–751 (2007).

28. Kendall, D.G. Shape manifolds, procrustean metrics, and complex projective spaces. Bulletin of the London mathematical society 16, 81–121 (1984).

29. Gower, J.C. Generalized procrustes analysis. Psychometrika 40, 33–51 (1975).

30. Rohlf, F.J. & Slice, D. Extensions of the Procrustes Method for the Optimal Superimposition of Landmarks. Syst Zool 39, 40–59 (1990).

31. O’Higgins, P., Chadfield, P. & Jones, N. Facial growth and the ontogeny of morphological variation within and between the primates Cebus apella and Cercocebus torquatus. Journal of Zoology 254, 337–357 (2001).

32. Dryden, I.L. & Mardia, K.V. Multivariate shape analysis. Sankhya 55, 460–480 (1993).

33. Young, M.D. & Behjati, S. SoupX removes ambient RNA contamination from droplet-based single-cell RNA sequencing data. Gigascience 9 (2020).

34. Subramanian, A., Kuehn, H., Gould, J., Tamayo, P. & Mesirov, J.P. GSEA-P: a desktop application for Gene Set Enrichment Analysis. Bioinformatics 23, 3251–3253 (2007).

35. Huang da, W., Sherman, B.T. & Lempicki, R.A. Systematic and integrative analysis of large gene lists using DAVID bioinformatics resources. Nature protocols 4, 44–57 (2009).

36. Huang da, W., Sherman, B.T. & Lempicki, R.A. Bioinformatics enrichment tools: paths toward the comprehensive functional analysis of large gene lists. Nucleic acids research 37, 1–13 (2009).

37. Moh, A. et al. Role of STAT3 in liver regeneration: survival, DNA synthesis, inflammatory reaction and liver mass recovery. Laboratory investigation; a journal of technical methods and pathology 87, 1018–1028 (2007).

38. Rashid, H., Chen, H., Hassan, Q. & Javed, A. Dwarfism in homozygous Agc1. Genesis 55 (2017).

39. Alberton, P. et al. Aggrecan Hypomorphism Compromises Articular Cartilage Biomechanical Properties and Is Associated with Increased Incidence of Spontaneous Osteoarthritis. Int J Mol Sci 20 (2019).

40. Xiao, Z.-f. et al. Osteoporosis of the vertebra and osteochondral remodeling of the endplate causes intervertebral disc degeneration in ovariectomized mice. Arthritis Research & Therapy 20, 207 (2018).

41. Zieba, J. et al. TGFβ and BMP Dependent Cell Fate Changes Due to Loss of Filamin B Produces Disc Degeneration and Progressive Vertebral Fusions. PLoS Genet 12, e1005936 (2016).

42. Tsingas, M. et al. Sox9 deletion causes severe intervertebral disc degeneration characterized by apoptosis, matrix remodeling, and compartment-specific transcriptomic changes. Matrix Biol 94, 110–133 (2020).

43. Bookstein, F.L. Morphometric tools for landmark data: geometry and biology. (Cambridge University Press, 1997).

44. Marcus, L.F., Corti, M., Loy, A., Naylor, G.J. & Slice, D.E. Advances in morphometrics, Vol. 284. (Springer Science & Business Media, 2013).

45. O’Higgins, P. The study of morphological variation in the hominid fossil record: biology, landmarks and geometry. The Journal of Anatomy 197, 103–120 (2000).

46. Dryden, I.L. & Mardia, K.V. Multivariate shape analysis. Sankhyā: The Indian Journal of Statistics, Series A, 460–480 (1993).

47. Bookstein, F.L. in Biennial International Conference on Information Processing in Medical Imaging 326–342 (Springer, 1991).

48. Decker, R.S., et al. Cell origin, volume and arrangement are drivers of articular cartilage formation, morphogenesis and response to injury in mouse limbs. Developmental biology 426, 56–68 (2017).

49. Dreier, R. Hypertrophic differentiation of chondrocytes in osteoarthritis: the developmental aspect of degenerative joint disorders. Arthritis Research & Therapy 12, 216 (2010).

50. Bi, W., Deng, J.M., Zhang, Z., Behringer, R.R. & de Crombrugghe, B. Sox9 is required for cartilage formation. Nat Genet 22, 85–89 (1999).

51. Smits, P. et al. The transcription factors L-Sox5 and Sox6 are essential for cartilage formation. Dev Cell 1, 277–290 (2001).

52. Chan, H.Y. et al. Comparison of IRES and F2A-based locus-specific multicistronic expression in stable mouse lines. PLoS One 6, e28885 (2011).

53. Li, J. et al. Systematic Reconstruction of Molecular Cascades Regulating GP Development Using Single-Cell RNA-Seq. Cell Rep 15, 1467–1480 (2016).

54. Ai, G. et al. Epidermal growth factor promotes proliferation and maintains multipotency of continuous cultured adipose stem cells via activating STAT signal pathway in vitro. Medicine 96, e7607 (2017).

55. Li, D.-q., Wan, Q.-l., Pathak, J.L. & Li, Z.-b. Platelet-derived growth factor BB enhances osteoclast formation and osteoclast precursor cell chemotaxis. Journal of Bone and Mineral Metabolism 35, 355–365 (2017).

56. Barclay, J.L. et al. In vivo targeting of the growth hormone receptor (GHR) Box1 sequence demonstrates that the GHR does not signal exclusively through JAK2. Molecular endocrinology (Baltimore, Md.) 24, 204–217 (2010).

57. Sundberg, J.P. et al. Systematic screening for skin, hair, and nail abnormalities in a large-scale knockout mouse program. PLoS One 12, e0180682–e0180682 (2017).

58. Ware, C.B. et al. Targeted disruption of the low-affinity leukemia inhibitory factor receptor gene causes placental, skeletal, neural and metabolic defects and results in perinatal death. Development 121, 1283–1299 (1995).

59. Mikelonis, D., Jorcyk, C.L., Tawara, K. & Oxford, J.T. Stüve-Wiedemann syndrome: LIFR and associated cytokines in clinical course and etiology. Orphanet Journal of Rare Diseases 9, 34 (2014).

60. Grumbach, M.M. Estrogen, bone, growth and sex: a sea change in conventional wisdom. J Pediatr Endocrinol Metab 13 Suppl 6, 1439–1455 (2000).

61. Henry, S.P., Liang, S., Akdemir, K.C. & de Crombrugghe, B. The postnatal role of Sox9 in cartilage. J Bone Miner Res 27, 2511–2525 (2012).

62. Sims, N.A. et al. Glycoprotein 130 regulates bone turnover and bone size by distinct downstream signaling pathways. J Clin Invest 113, 379–389 (2004).

63. Wang, M. et al. Both endogenous and exogenous testosterone decrease myocardial STAT3 activation and SOCS3 expression after acute ischemia and reperfusion. Surgery 146, 138–144 (2009).

64. Dziennis, S., Jia, T., Rønnekleiv, O.K., Hurn, P.D. & Alkayed, N.J. Role of signal transducer and activator of transcription-3 in estradiol-mediated neuroprotection. Journal of Neuroscience 27, 7268–7274 (2007).

65. Ahima, R.S., Dushay, J., Flier, S.N., Prabakaran, D. & Flier, J.S. Leptin accelerates the onset of puberty in normal female mice. J Clin Invest 99, 391–395 (1997).

66. Di Domenico, F. et al. Involvement of stat3 in mouse brain development and sexual dimorphism: A proteomics approach. Brain Research 1362, 1–12 (2010).

67. Wegrzyn, J. et al. Function of mitochondrial Stat3 in cellular respiration. Science 323, 793–797 (2009).

68. Flores, A. et al. Lactate dehydrogenase activity drives hair follicle stem cell activation. Nature cell biology 19, 1017–1026 (2017).

69. Lichanska, A.M. & Waters, M.J. How growth hormone controls growth, obesity and sexual dimorphism. Trends in Genetics 24, 41–47 (2008).

70. Shim, K.S. Pubertal growth and epiphyseal fusion. Ann Pediatr Endocrinol Metab 20, 8–12 (2015).

71. Rhee, N., Jeong, K., Yang, E.M. & Kim, C.J. Gigantism caused by growth hormone secreting pituitary adenoma. Ann Pediatr Endocrinol Metab 19, 96–99 (2014).

72. Kant, S.G. et al. A novel variant of FGFR3 causes proportionate short stature. Eur J Endocrinol 172, 763–770 (2015).

73. Chen, Y.H. et al. Absence of GP130 cytokine receptor signaling causes extended Stuve-Wiedemann syndrome. J Exp Med 217 (2020).

74. Sederquist, B., Fernandez-Vojvodich, P., Zaman, F. & Sävendahl, L. RECENT RESEARCH ON THE GROWTH PLATE: Impact of inflammatory cytokines on longitudinal bone growth. 53, T35 (2014).

75. Jikko, A. et al. Effects of interleukin-6 on proliferation and proteoglycan metabolism in articular chondrocyte cultures. Cell biology international 22, 615–621 (1998).

76. Nasi, S., So, A., Combes, C., Daudon, M. & Busso, N. Interleukin-6 and chondrocyte mineralisation act in tandem to promote experimental osteoarthritis. Annals of the rheumatic diseases 75, 1372–1379 (2016).

77. Legendre, F., Dudhia, J., Pujol, J.-P. & Bogdanowicz, P. JAK/STAT but not ERK1/ERK2 pathway mediates interleukin (IL)-6/soluble IL-6R down-regulation of type II collagen, aggrecan core, and link protein transcription in articular chondrocytes association with a down-regulation of SOX9 expression. Journal of Biological Chemistry 278, 2903–2912 (2003).

78. Latourte, A. et al. Systemic inhibition of IL-6/Stat3 signalling protects against experimental osteoarthritis. Annals of the Rheumatic Diseases 76, 748–755 (2017).

79. Ryu, J.H. et al. Interleukin-6 plays an essential role in hypoxia-inducible factor 2α–induced experimental osteoarthritic cartilage destruction in mice. Arthritis & Rheumatism 63, 2732–2743 (2011).

80. Bharucha, K.N. et al. Growth During Tocilizumab Therapy for Polyarticular-course Juvenile Idiopathic Arthritis: 2-year Data from a Phase III Clinical Trial. The Journal of Rheumatology 45, 1173 (2018).

